# Protein kinase D1 phosphorylates CBX8 to facilitate the disassociation of PRC1 complex from p16 promoter and promotes cell senescence

**DOI:** 10.1101/2020.08.09.242891

**Authors:** Yuanyuan Su, Yao Liang, Chenzhong Xu, Na Zhang, Doudou Liu, Guodong Li, Tanjun Tong, Jun Chen

## Abstract

The Polycomb group (PcG) protein chromobox 8 (CBX8) is the subunit of Polycomb repressive complex 1 (PRC1) and recognizes the trimethylation of histone H3 on Lysine 27 (H3K27me3), and coordinates with PRC2 complex to function as epigenetic gene silencer. CBX8 plays a key role in cell proliferation, stem cell biology, cell senescence, and cancer development. However, our knowledge of CBX8 post-translational modifications remains elusive. Here, we report that protein kinase D1 (PKD1) interacts and phosphorylates CBX8 at Ser256 and Ser311 in an evolutionarily conserved motif. We found that PKD1 activation triggered by serum stimulation, Nocodazole treatment and oncogene Ras-induced cell senescence (Ras OIS) all promotes CBX8 S256/311 phosphorylation. PKD1-mediated CBX8 S256/311 phosphorylation impairs PRC1 complex integrity by reducing the binding of CBX8 to other PRC1 components BMI1 and RING1B, decreases the monoubiquitination of histone H2AK119, and results in CBX8 dissociation from its target INK4a/ARF locus and the de-repression of p16, and thus ultimately facilitates cellular senescence. CBX8 S256/311 phosphorylation also compromises hepatocellular cancer cells proliferation and migration. Collectively, these results suggest that PKD1-mediated CBX8 S256/311 phosphorylation is a key mechanism governing CBX8 function, including cell senescence and cancer cell proliferation.

**Financial support:** This work was supported by grants from Ministry of Science and Technology of the People’s Republic of China (2018YFC2000102), and from National Natural Science Foundation of China (31871382 and 81571369).

## Introduction

Cellular senescence is a permanent cell cycle arrest state which can be triggered by diverse intracellular and extracellular stimuli, such as telomere shortening, DNA damage, oxidative stress, oncogenes activation and tumor suppressor genes loss, etc. These stimuli are signaled through activating two major tumor suppressor pathways p53/p21^WAF1^ and p16^INK4a^/pRB to inhibit cell cycle irreversibly(Salama et al., 2014). Cellular senescence is recognized as a bona fide tumor-suppressive mechanism in vivo, however, it also contributes to aging and many age-related diseases including cancer in late-life(He and Sharpless, 2017; Munoz-Espin and Serrano, 2014).

The cyclin-dependent kinase (CDK) inhibitor p16^INK4a^ is encoded by INK4a/ARF locus (CDKN2A), which also encodes p14^ARF^ (or p19^Arf^ in mice). In most primary cells, the INK4a/ARF locus is tightly regulated by the Polycomb group (PcG) proteins that function as epigenetic modifiers and transcriptional repressors(Gil and Peters, 2006). PcG proteins form two distinct protein complexes called Polycomb Repressive Complex 1 (PRC1) and 2 (PRC2). In mammals, PRC2 composes four core proteins EZH1/2, EED, SUZ12 and RbAp 48. PRC1 consists of four core subunits including polycomb group finger proteins PCGF1–6 (such as MEL18 and BMI1), chromobox (CBX) proteins (including CBX2, 4, 6, 7, 8), PHC proteins PHC1–3, and RING proteins (including RING1A/B)(Chittock et al., 2017; Connelly and Dykhuizen, 2017). PRC2 component EZH1/2 is a histone lysine methyltransferase that catalyzes the trimethylation of histone H3 lysine 27 (H3K27me3) primarily at promoters of target genes(Cao et al., 2002). This transcriptionally repressive histone mark is then recognized by CBX proteins containing chromodomain in PRC1, where the histone H2A at Lys119 is mono-ubiquitinated (H2AK119ub1) by the E3 ubiquitin ligase RING1A/B in PRC1(Cao et al., 2005; Wang et al., 2004). The presence of both PRC2 and PRC1 in the promoter region acts in concert to facilitate epigenetic silencing of target genes(Aranda et al., 2015). PcGs play key roles in development, stem cell maintenance, senescence, and cancer(Entrevan et al., 2016; Piunti and Shilatifard, 2016). One of the critical PcGs-repressed targets is the INK4a/ARF locus(Gil and Peters, 2006). During cell senescence, many PcG proteins including CBX8 are down-regulated, leading to the loss of H3K27me3 at the INK4a/ARF locus, which results in the upregulation of p16(Bracken et al., 2007; Dietrich et al., 2007; Gil et al., 2004; Jacobs et al., 1999). On the other hand, ectopic expression of CBX8 or other PcG proteins delay or bypass the onset of senescence in mouse and human fibroblasts.

Growing evidences suggest that PcG proteins are subjected to the post-translational modifications, particularly phosphorylation, in response to the environment or cell signaling cues(Chen et al., 2010; Kaneko et al., 2010; Kawaguchi et al., 2017; Liu et al., 2012; Voncken et al., 2005; Wan et al., 2018; Wei et al., 2011; Wu et al., 2013; Wu and Zhang, 2011). EZH2 can be phosphorylated by CDK1, Akt and AMP-activated protein kinase (AMPK), leading to the alterations of PRC complexes stability and enzymatic activity, thereby affecting target genes transcription(Chen et al., 2010; Kaneko et al., 2010; Liu et al., 2012; Wan et al., 2018; Wei et al., 2011; Wu and Zhang, 2011). CBX7 is phosphorylated by mitogen-activated protein kinase (MAPK) to enhance PRC1 association and target genes silencing(Wu et al., 2013). It is reported that CBX2 phosphorylation controls its nucleosome-binding specificity(Kawaguchi et al., 2017). Although the phosphorylation sites of CBX8 have been identified by global phosphoproteomic analyses, however, these sites have never been validated in vivo and the relevant kinases and functions remain unknown(Zhou et al., 2013). PIM1 is reported to phosphorylate CBX8 to promote its degradation, thereby up-regulating p16 to enhance cell senescence, however, the phosphorylation site has not been identified in this study(Zhan et al., 2018).

Protein kinase D (PKD) is a serine/threonine kinase family belonging to the calcium/calmodulin-dependent kinase (CAMK) superfamily, and comprises PKD1, PKD2, and PKD3 isoforms(Fu and Rubin, 2011; Rozengurt, 2011). PKD1, the best characterized PKD, regulates multiple biological processes and pathologies, such as oxidative stress response(Storz et al., 2005), cell adhesion and motility(Eiseler et al., 2009), protein transport from trans-Golgi network to cell surface(Hausser et al., 2005; Malhotra and Campelo, 2011), immune response(Haxhinasto and Bishop, 2003), and cancer, etc.(Roy et al., 2017). Particularly, PKD1 can phosphorylate class IIa histone deacetylases (HDACs) HDAC5 and HDAC7 in the nucleus, leading them nuclear export, thereby releasing HDAC5/7-repressed genes transcription to regulate diverse biological functions(Fielitz et al., 2008; Jensen et al., 2009; Vega et al., 2004; Wang et al., 2008; Xu et al., 2007). We previously report that PKD1 activates both NF-κB and classical protein secretory pathway to modulate senescence-associated inflammatory cytokines interleukin-6 (IL-6) and interleukin-8 (IL-8) expressions and secretion thereby promoting cellular senescence(Su et al., 2018; Wang et al., 2014). However, whether PKD1 can regulate cellular senescence at epigenetic level remains unknown.

Here, we report that PKD1 directly interacts with CBX8 and phosphorylates CBX8 at Ser256 and Ser311, which is observed in oncogene Ras-induced senescence (Ras OIS), serum stimulation and Nocodazole treatment. Functionally, CBX8 phosphorylation by PKD1 enhances p16 expression and cell senescence, and suppresses liver cancer cell proliferation and migration. Mechanistically, CBX8 phosphorylation attenuates PRC1 complex association and H2AK119ub1 level, but has no impact on H3K27me3 level. Collectively, we report the novel CBX8 phosphorylation sites and demonstrate that PKD1-mediated CBX8 phosphorylation at S256/S311 regulates PRC1 function.

## Results

### PKD1 directly interacts with CBX8 and phosphorylates CBX8

To better understand biological processes regulated by PKD1, we attempted to identify the interactors and potential novel substrates of PKD1. To this end, FLAG-PKD1 was ectopically expressed in HEK293T cells and immunoprecipitated by using Flag antibody. The eluted proteins were separated and silver stained, and then subjected to LC–MS/MS analysis (Fig. 1A). Mass spectrometric analysis indicated that CBX8 and RING1B, both of them belonging to the PRC1 complex, were in the presence of the PKD1 interactome (Fig. 1A).

**Figure 1.**
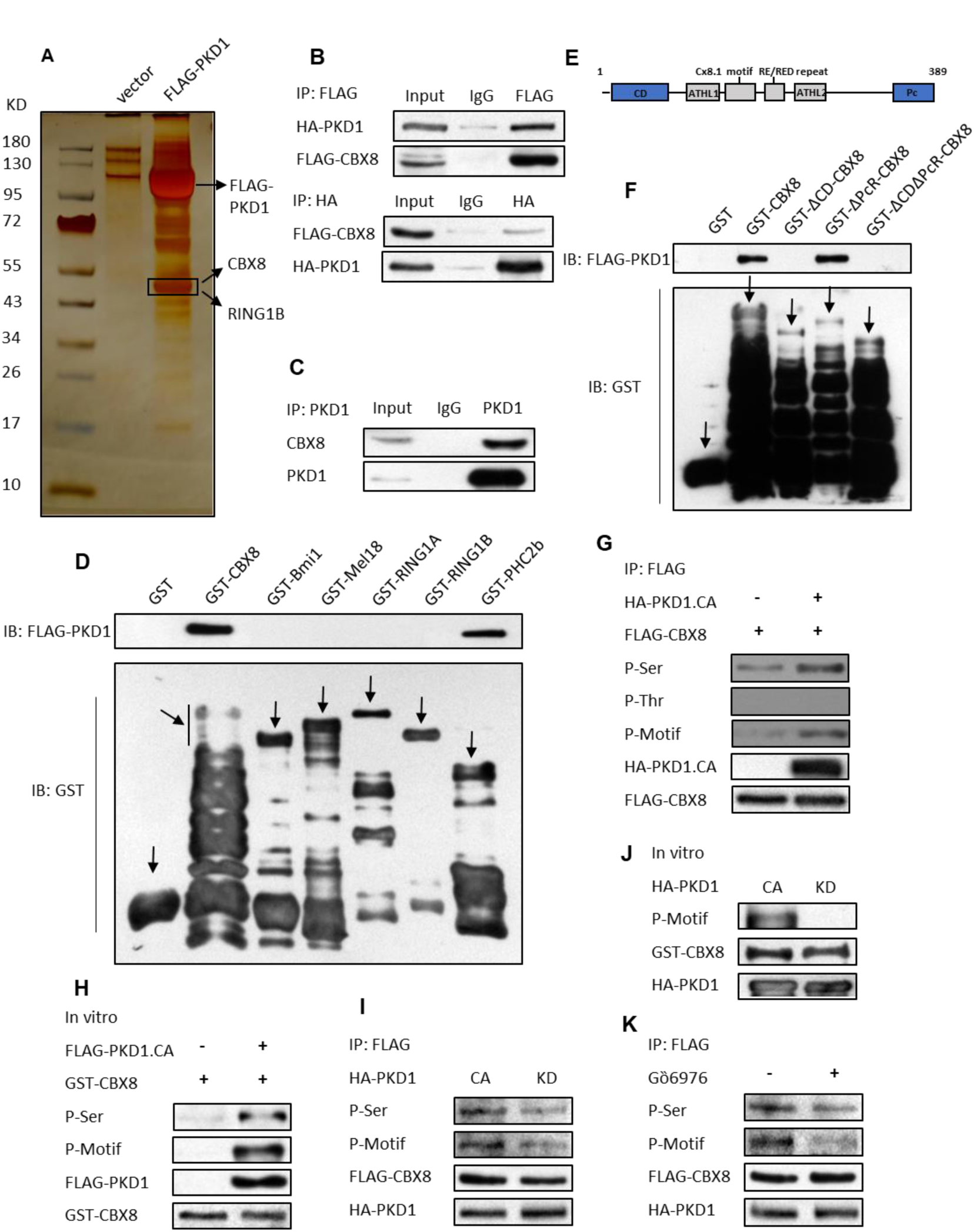
PKD1 associates with CBX8 and phosphorylates CBX8. (A) Mass spectrometry analysis of PKD1-associated proteins. Cellular extracts from HEK293T cells overexpressing FLAG-PKD1 were immunoprecipitated with anti-FLAG antibody and Protein G-Sepharose beads. After extensively washing the beads, immobilized proteins were eluted in SDS–PAGE sample buffer. The eluent was resolved by SDS–PAGE and silver stained. The protein bands were retrieved and analyzed by mass spectrometry. (B) HEK 293T cells were transfected with FLAG-CBX8 and HA-PKD1 plasmids. IP assay and subsequent western blot analysis were performed by using the indicated antibodies. (C) Whole-cell lysates from HEK293T cells were immunoprecipitated with anti-PKD1 antibody followed by immunoblotting with the indicated antibodies. (D) GST pull-down experiments were performed with bacterially expressed GST-tagged proteins and HEK293T-expressed FLAG-PKD1. (E) The schematic representation of the key domains within CBX8 was depicted. The chromodomain and Pc box are enclosed in a blue dotted box. (F) GST pull-down assay was performed by incubating GST-CBX8 or its various deletion mutants with HEK293T-expressed FLAG-PKD1. (G) HEK293T cells were co-transfected FLAG-CBX8 with or without HA-PKD1.CA. An IP assay was carried out using anti-FLAG antibody followed by western blotting with anti-HA, anti-FLAG, anti-pMOTIF, anti-phosphothreonine, anti-phosphoserine antibodies. (H) Purified GST-CBX8 protein was incubated with or without FLAG-PKD1.CA in the presence of ATP. The reaction products were separated by SDS-PAGE and immunoblotted with the indicated antibodies. (I) HEK293T cells were co-transfected FLAG-CBX8 with HA-PKD1.CA or HA-PKD1.KD. An IP assay was carried out using anti-FLAG antibody followed by immunoblotting with the indicated antibodies. (J) Purified GST-CBX8 protein was incubated with FLAG-PKD1.CA or FLAG-PKD1.KD in the presence of ATP. The reaction products were separated by SDS-PAGE and immunoblotted with the indicated antibodies. (K) HEK293T cells were transfected with FLAG-CBX8 and treated with or without Gö6976. An IP assay was carried out using anti-FLAG antibody followed by immunoblotting with the indicated antibodies.

We then validated the interaction between PKD1 and CBX8 by using reciprocal co-immunoprecipitation (Co-IP) assays with anti-Flag or anti-HA antibodies, which demonstrated an association between HA-tagged PKD1 and FLAG-tagged CBX8 (Fig. 1B). Importantly, this association was also detected with endogenous PKD1 and CBX8 by Co-IP, indicating that these two molecules interact with each other in vivo (Fig. 1C). We further confirmed that PKD1 directly binds to CBX8 in vitro using the GST-pulldown assay (Fig. 1D). The direct association between PKD1 and PRC1 complex component PHC2b was also observed. However, no direct interaction was detected between PKD1 and other PRC1 complex components including RING1B (Fig. 1D). CBX8 protein possesses an N-terminal chromodomain (CD) and a C-terminal Polycomb Repressor box (PcR) (Fig. 1E)(Chittock et al., 2017; Connelly and Dykhuizen, 2017). To map the region of CBX8 responsible for binding to PKD1, we generated a series of GST-tagged CBX8 truncated mutants deleting either CD domain (GST-∆CD CBX8) or PcR domain (GST-∆PcR CBX8), or both of them (GST-∆CD∆PcR CBX8), respectively. GST-pulldown results showed that the binding ability of CBX8-ΔCD mutant to PKD1 completely lost, whereas CBX8-∆PcR mutant preserved the similar binding ability to PKD1, when compared with wild-type CBX8 (WT-CBX8) (Fig. 1F). This result indicates that the CD domain of CBX8 is critical for CBX8 binding to PKD1.

Since PKD1 is a serine/threonine kinase, we next investigated whether PKD1 phosphorylates CBX8. To this end, FLAG-CBX8 was transfected with or without constitutive active PKD1 mutant HA-PKD1-CA (PKD1-S738/742E double mutations) in HEK293T cells, then followed by immunoprecipitation of FLAG-CBX8 and subjected to immunoblotting analysis using PKD1 substrate motif antibody (p-Motif antibody) which recognizes PKD1-phosphorylated substrates at its consensus motif as well as antibodies recognizing phospho-serine residue or phospho-threonine residue, to detect CBX8 phosphorylation status in the PKD1-CA-overexpressing cells. The results showed that the exogenously expressed FLAG-CBX8 could be phosphorylated by PKD1-CA in cells and CBX8 was serine-phosphorylated but not threonine-phosphorylated (Fig. 1G). We also investigated CBX8 phosphorylation in vitro by incubation purified GST-tagged CBX8 with or without purified FLAG-PKD1-CA in the presence of adenosine triphosphate (ATP). The result demonstrated that GST-CBX8 was efficiently serine-phosphorylated by PKD1-CA in in vitro kinase assay (Fig. 1H). In contrast, the kinase-inactive PKD1 mutant PKD1-KD (PKD1-K612W mutation) was unable to phosphorylate CBX8 in vitro and in vivo compared to PKD1-CA (Fig. 1I and J). Similarly, PKD1 inhibitor Gö6976 treatment also suppressed PKD1-CA-mediated CBX8 phosphorylation at serine residue (Fig. 1K). It was worth to mention that although PKD1 could associate with PHC2b, it did not phosphorylate PHC2b (Fig. S1A). Furthermore, the possibility that PKD1 phosphorylating other PRC1 core components was excluded by the result that GST-CBX8 was the only target phosphorylated by PKD1-CA, but not other PRC1 core components including BMI1, MEL18, and RING1A/B (Fig. S1B). Taken together, these results demonstrate that PKD1 directly interacts with CBX8 and phosphorylates CBX8 at serine residue.

### PKD1 phosphorylates CBX8 at Ser256 and Ser311

To determine which serine residue of CBX8 may be phosphorylated by PKD1, we utilized a phosphorylation site prediction program MotifScan. We found that CBX8 harbored two putative phosphorylation sites at Ser256 and Ser311 that perfectly matched PKD1 phosphorylation consensus motif (L/V/IXRXXS/T, where X is any residue). These two residues are highly conserved in humans, mice, and rats (Fig. 2A). We then performed in vitro phosphorylation of CBX8 and used mass spectrometry to identify which residue of CBX8 was phosphorylated by PKD1-CA. Three phosphorylated serine sites including Ser219, Ser236, and Ser256 were mapped by mass spectrometry (Supplemental Table 1).

**Figure 2.**
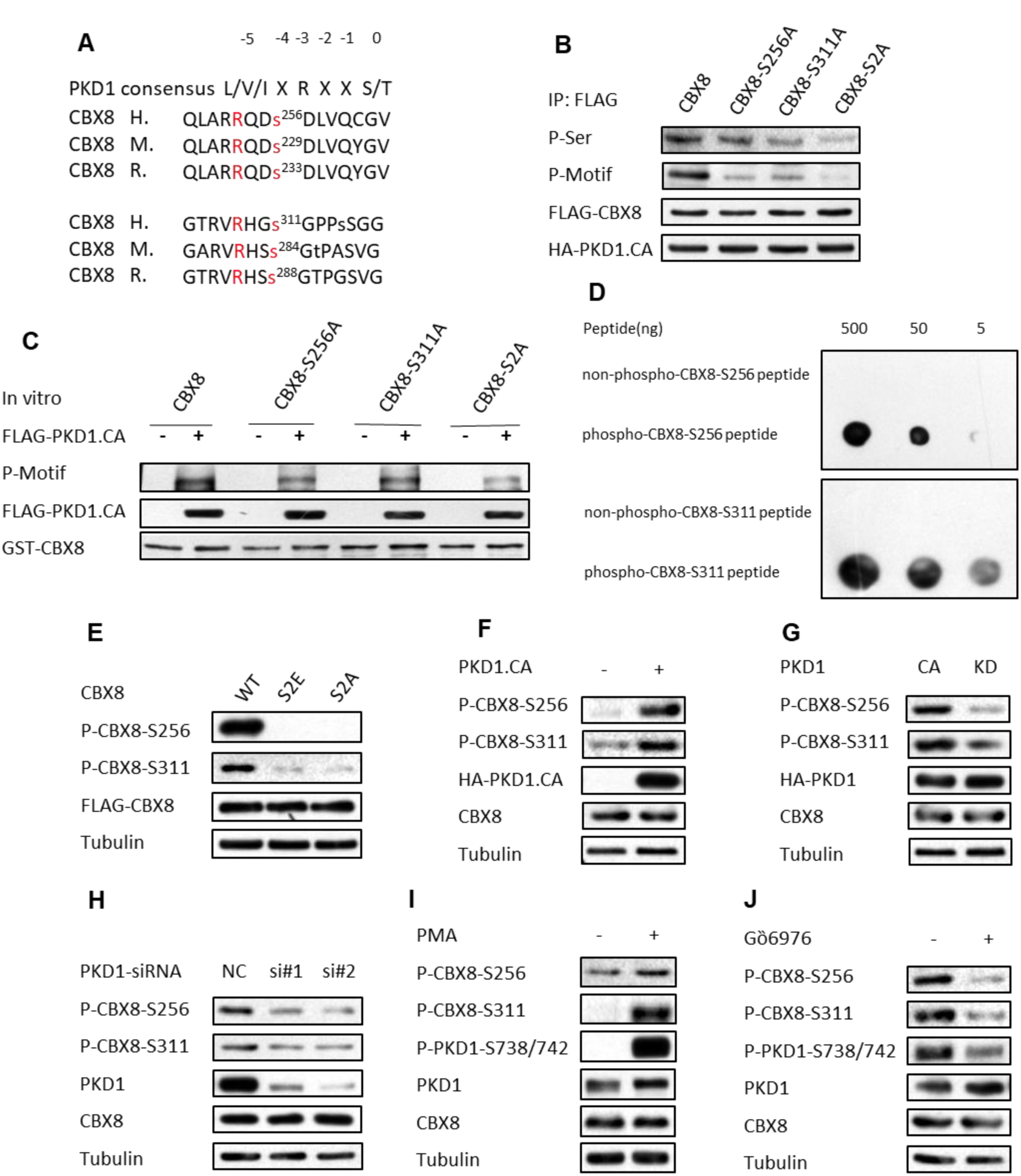
PKD1 phosphorylates CBX8 at Ser256 and Ser311. (A) Ser256 and Ser311 residues of CBX8 lie within the PKD1 phosphorylation consensus motif. These two sites are conserved through species: H. sapiens, Homo sapiens; M. musculus, Mus musculus; R. norvegicus, Rattus norvegicus. (B) HEK293T cells were co-transfected WT-CBX8 or its mutants with or without HA-PKD1.CA. An IP assay was carried out using anti-FLAG followed by immunoblotting with the indicated antibodies. (C) Purified GST-tagged WT-CBX8 or its mutants were incubated with or without FLAG-PKD1.CA in the presence of ATP. The reaction products were separated by SDS-PAGE and immunoblotted with the indicated antibodies. (D) Dot blot assay was performed using p-CBX8-S256 and p-CBX8-S311 antibodies to detect CBX8 peptides containing phosphorylated or non-phosphorylated Ser256 or Ser311. (E) HEK293T cells transfected with FLAG-CBX8 or its various mutants, then the phosphorylation levels of CBX8 at S256 or S311 were detected by western blot analysis using p-CBX8-S256 and p-CBX8-S311 antibodies. (F-H) HEK293T cells transfected with or without HA-PKD1.CA (F), with HA-PKD1.CA or PKD1.KD (J), with NC or PKD1-siRNA#1, #2 (H), were performed western blot using the indicated antibodies. (I and J) HEK293T cells treated with Gö6976 or PMA were assessed by western blot using the indicated antibodies.

To validate whether PKD1 phosphorylates CBX8 at these sites, we mutated these four serine residues to alanine individually or simultaneously to generate five mutants: CBX8-S219A, S236A, S256A, S311A, and Ser256/311A. The PKD1-mediated phosphorylation levels of CBX8-S219A and S236A mutants were comparable to WT-CBX8 phosphorylation level (Fig. S2), indicating that these two serine residues are not the major phosphorylation sites in CBX8 by PKD1. Either CBX8-S256A or S311A mutant displayed a moderate reduction in PKD1-mediated serine phosphorylation level both in cells and in vitro kinase assay, whereas the double-residues mutant CBX8-S256/311A showed a marked reduction in serine phosphorylation level, when compared with WT-CBX8 phosphorylation level (Fig. 2B and C). Collectively, these results suggest that Ser256 and Ser311 are two major sites in CBX8 phosphorylated by PKD1 in vivo.

To further confirm that PKD1 does indeed phosphorylate CBX8 at Ser256/311 in vivo, we generated two antibodies that recognizes CBX8 phosphorylation at Ser256 or Ser311 (p-CBX8-S256 and p-CBX8-S311), respectively. Dot blot analysis validated that these two antibodies specifically recognized the peptides containing the phosphorylated Ser256 or Ser311 residues of CBX8, but failed to detect the unphosphorylated CBX8 peptides (Fig. 2D). The specificities of these two antibodies were further proved by that p-CBX8-S256 and p-CBX8-S311antibodies only recognized exogenously expressed WT-CBX8, but not CBX8 mutants which Ser256 and Ser311 were mutated to alanine (CBX8-S256/311A, S2A) or glutamic acid (CBX8-S256/311E, S2E) (Fig. 2E).

We then used these two antibodies to determine whether PKD1 phosphorylates CBX8 at Ser256/311 in vivo. Ectopic expression of PKD1-CA largely increased endogenous WT-CBX8 phosphorylation levels at Ser256 and Ser311 in HEK293T cells (Fig. 2F), whereas PKD1-KD could not promote CBX8 phosphorylation (Fig. 2G). Furthermore, PKD1 knockdown by PKD1-siRNAs repressed CBX8 phosphorylation (Fig. 2H), suggesting that PKD1 is required for CBX8 phosphorylation at Ser256/311. Similar to PKD1-CA effect, PKD1 activation by PMA treatment also markedly induced endogenous CBX8 phosphorylation at Ser256/311 (Fig. 2I). In contrast, PKD1 inhibition by Gö6976 treatment suppressed endogenous CBX8 phosphorylation at Ser256/311 (Fig. 2J). Collectively, these results indicate that PKD1 phosphorylates CBX8 at both Ser256 and Ser311 residues in vitro and in vivo.

### Activation of PKD1 by serum stimulation or Nocodazole treatment promotes CBX8 S256/311 phosphorylation

PKD1 is activated by mitogenic signals, such as serum growth factors(Zugaza et al., 1996). Using these two specific antibodies, we examined whether activation of PKD1 by serum stimulation could induce CBX8 S256/311 phosphorylation. To this end, FLAG-CBX8 was transfected into H1299 and HEK293T cells followed by serum starvation, then serum was reintroduced and CBX8 S256/311 phosphorylation was monitored. Upon serum starvation, CBX8 S256/311 phosphorylation and PKD1 S738/742 phosphorylation were undetectable or barely detected. However, CBX8 S256/311 phosphorylation levels markedly increased and reached to peak levels within 1 to 2 h of serum reintroduction (Fig. 3A, and Fig. S3A). The CBX8 phosphorylation was sustained for up to 3-4 h after serum stimulation. PKD1 was also phosphorylated at S738/742 after serum reintroduction. Importantly, PKD1 activation correlated well with CBX8 phosphorylation (Fig. 3A, and Fig. S3A). Moreover, we found that knockdown of endogenous PKD1 by PKD1-siRNAs or inhibition of PKD1 activity by its inhibitor Gö6976 drastically suppressed endogenous CBX8 S256/311 phosphorylation induced by serum when compared with the corresponding control cells (Fig. 3B, C, and Fig. S3B, C). Altogether, these results suggest that PKD1 phosphorylates CBX8 at S256/311 during serum stimulation.

**Figure 3.**
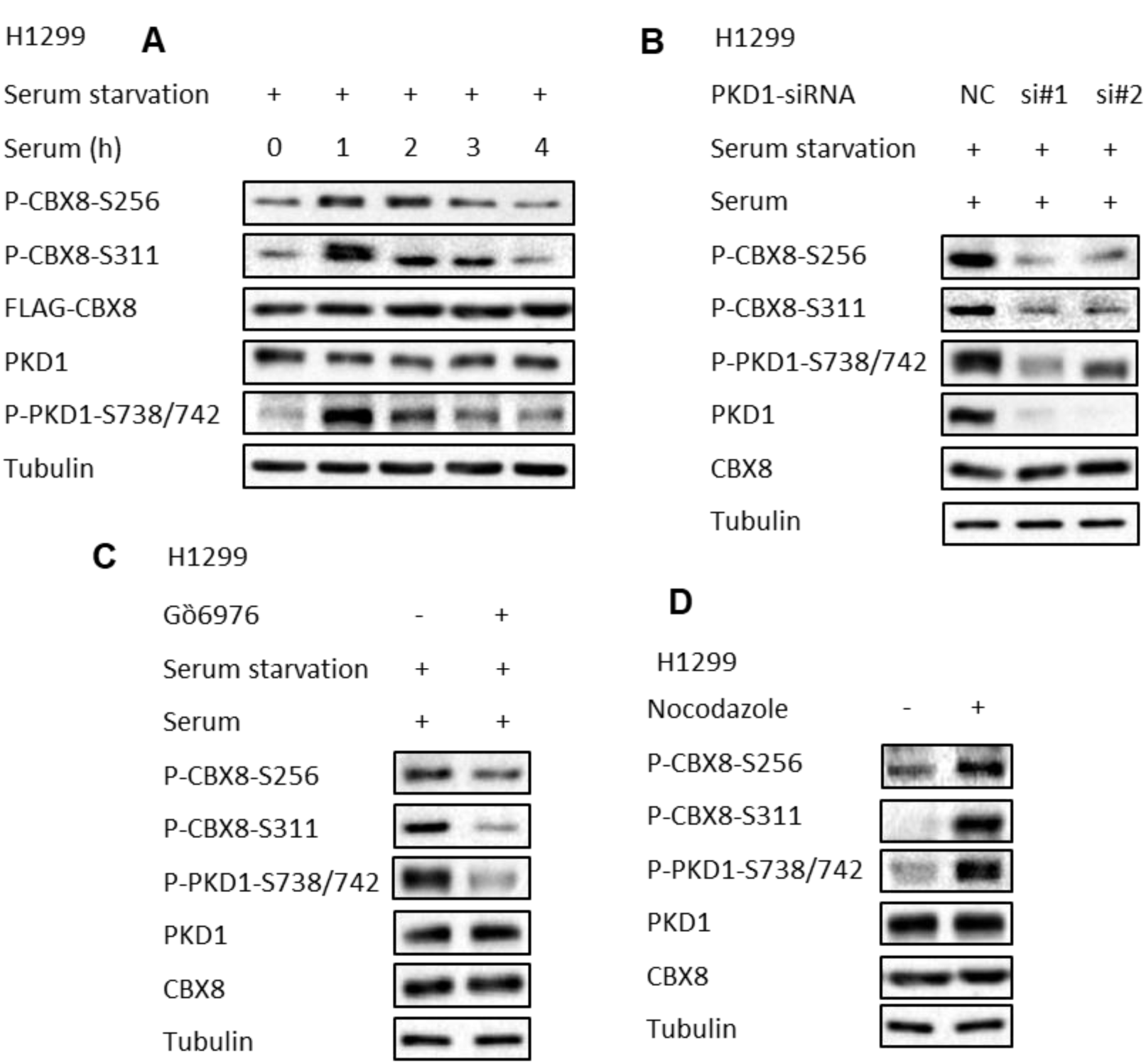
Serum stimulation or Nocodazole treatment-induced PKD1 activation promotes CBX8 S256/311 phosphorylation. (A) H1299 cells were deprived of serum for 24h, then serum was reintroduced in the culture for the indicated times, and western blot was performed using the indicated antibodies. (B) H1299 cells were transfected with PKD1-siRNAs for 48 h, then serum was deprived for 24h. Cells were then added serum for 1h and harvested for immunoblotting with the indicated antibodies. (C) H1299 cells were deprived of serum for 24h with or without Gȍ6976 treatment, then serum was reintroduced for 1h, and western blot was performed using the indicated antibodies. (D) H1299 cells were treated with or without Nocodazole (100 ng/mL) for 16h, then cells were subjected to immunoblotting with the indicated antibodies.

PKD1 is also activated by Nocodazole treatment(Fuchs et al., 2009). Therefore, we further determined whether activation of PKD1 by Nocodazole treatment could induce CBX8 S256/311 phosphorylation. Compared to the untreated cells, Nocodazole treatment remarkably induced CBX8 S256/311 phosphorylation and PKD1 activation both in H1299 and HEK293T cells (Fig. 3D, and Fig. S3D). Taken together, these results provide a direct support that PKD1 is a physiological upstream kinase of CBX8 which phosphorylates CBX8 at S256/311.

### PKD1-mediated phosphorylation of CBX8 at S256/311 attenuates CBX8 cell senescence suppressive function

As ectopic expression of CBX8 is known to bypass cell senescence by directly suppressing the INK4a/ARF locus(Dietrich et al., 2007), and we previously also reveal that PKD1 promotes Ras OIS(Su et al., 2018; Wang et al., 2014), we then investigated whether PKD1 could phosphorylate CBX8 at S256/311 in Ras OIS and the biological importance of CBX8 S256/311 phosphorylation in cell senescence. To this end, we used IMR90 cells ectopically expressing oncogene Ras protein as a Ras OIS model, and the human lung fibroblast 2BS cells as a replicative senescence model. We first confirmed that co-expressed FLAG-CBX8 and HA-PKD1 also interacted with each other in both IMR90 and 2BS fibroblasts (Fig. S4A). Then we detected the phosphorylation levels of CBX8 at Ser256/311 during Ras OIS establishment course. Indeed, we observed a time-dependent increased CBX8 S256/311 phosphorylation upon Ras expression in IMR90 cells (Fig. 4A). Consistent with our previous result(Wang et al., 2014), PKD1 was gradually activated during Ras OIS. Importantly, the activation of PKD1 correlated well with CBX8 S256/311 phosphorylation. The senescent state of Ras OIS was characterized by the up-regulation of p16 and p21 (Fig. 4A).

**Figure 4.**
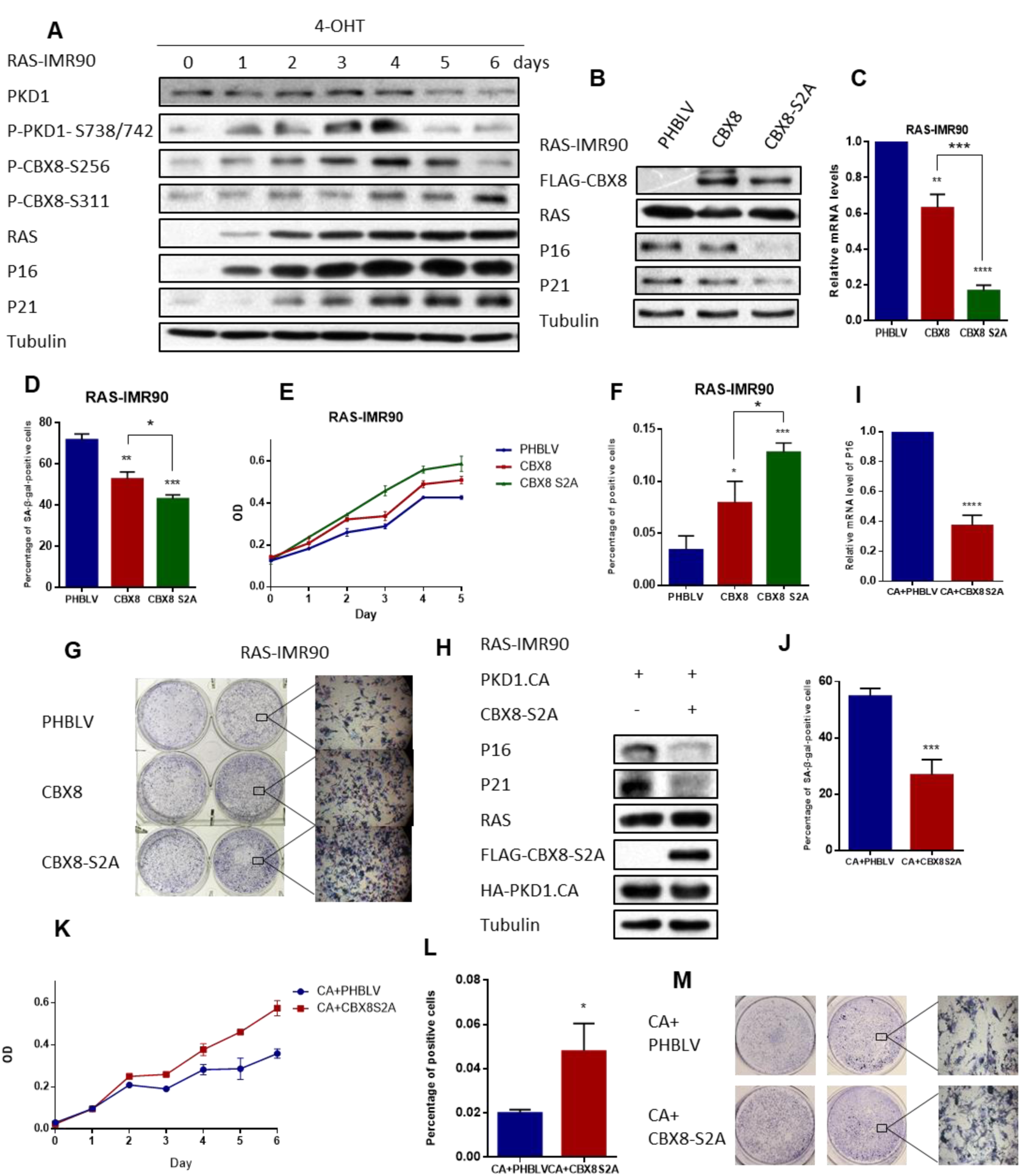
CBX8 S256/311 phosphorylation by PKD1 mitigates CBX8 cell senescence suppressive function. (A) ER: RAS IMR90 cells were induced senescent by 4-OHT treatment for 6 days, and then western blot was performed using the indicated antibodies. (B) PHBLV vector, FLAG-WT-CBX8 or FLAG-CBX8-S2A mutant were expressed in RAS-IMR90 cells with a lentiviral expression system. Then cells were subjected to western blot for the indicated proteins. (C) Total RNAs were extracted from (B) cells, then p16 mRNA level was measured by qPCR. (D) RAS-IMR90 cells infected with the indicated vectors were subjected to SA-β-gal activity staining. The percentages of SA-β-gal staining positive cells were statistically analyzed. Data are presented as mean ± s.d. from three independent experiments with 300 cells per experiment. (E) Growth curves of RAS-IMR90 cells infected with the indicated vectors were determined by CCK-8 assay. Data are presented as mean ± s.d. from three independent experiments, each performed in triplicate. (F) RAS-IMR90 cells infected with the indicated vectors were subjected to EdU incorporation assay. The percentages of EdU incorporating positive cells were statistically analyzed. Data are presented as mean ± s.d. from three independent experiments with 300 cells per experiment. (G) RAS-IMR90 cells infected with the indicated vectors were cultured for 14 days, then colony formation assay was performed by crystal violet staining cells. The representative magnified images (x40) were shown. (H-M) RAS-IMR90 cells were infected with HA-PKD1.CA lentiviral vector, then transfected with or without CBX8-S2A. Then cells were subjected to western blot (H), Realtime qPCR (I), SA-β-gal activity staining (J), cell growth assay (K), EdU incorporation assay (L), and colony formation assay (M), to evaluate cell senescent status.

We next exploited the biological function of CBX8 S256/311 phosphorylation in cell senescence. To this end, WT-CBX8, or unphosphorylatable mutant CBX8-S2A, was co-infected with RAS in young IMR90 cells, respectively. After 6 days Ras expression, cells were subjected to SA-β-gal activity assay, cell proliferation assay, EdU incorporation assay, and colony formation assay, respectively, to evaluate the cellular senescent state. As expected, compared to empty vector control, ectopic expression of WT-CBX8 substantially reduced p16 expressions at both protein and mRNA levels (Fig. 4B and C), decreased SA-β-gal-positive staining cells (Fig. 4D, and Fig. S4B), and increased cell proliferation and EdU incorporation as well as colony formation numbers (Fig. 4E-G, and Fig. S4C), indicating that CBX8 suppresses Ras OIS. Importantly, compared to WT-CBX8, CBX8-S2A mutant was more potent in repressing p16 expressions and SA-β-gal activity as well as in promoting cell proliferation and colony formation (Fig. 4B-G), suggesting that the unphosphorylatable mutant CBX-S2A may possess gain-of-function in suppressing Ras OIS.

We also stably overexpressed WT-CBX8, CBX8-S2A, and phospho-mimic mutant CBX8-S2E (Ser256/311 mutated to glutamic acid) in 50PD 2BS cells, respectively, to probe the role of CBX8 phosphorylation in replicative cell senescence. Similar to Ras OIS results, CBX8-S2A mutant was more potent than WT-CBX8 in attenuating 2BS cells replicative senescent phenotypes, including the decrease of p16 and SA-β-gal activity, and the elevation of cell proliferation, EdU incorporation and colony formation (Fig. S5A-H). In contrast, CBX8-S2E mutant was less potent than WT-CBX8 in mitigating 2BS cells replicative senescent phenotypes (Fig. S5I-M). Taken together, these results suggest that CBX8 phosphorylation at S256/311 by PKD1 facilitates cell senescence, which is supported by the observation that phospho-mimic mutant CBX8-S2E promotes cell senescence, whereas the unphosphorylatable mutant CBX8-S2A is more potent than WT-CBX8 in repressing cell senescence.

We previously show that PKD1 overexpression promotes Ras OIS, whereas PKD1 inhibition prevents Ras OIS(Storz et al., 2005). Therefore, we further explored whether PKD1 regulates Ras OIS partially via of PKD1-mediated CBX8 S256/311 phosphorylation. Compared to ectopic expression of PKD1-CA alone which is known to accelerate Ras OIS(Wang et al., 2008), the co-expression of CBX8-S2A with PKD1-CA antagonized the promoting effect of PKD1-CA on Ras OIS, which displayed the down-regulation of p16, p21 and SA-β-gal activity, and the increase of cell proliferation, EdU incorporation and colony formation (Fig. 4H-M, and Fig. S6A and B). Meanwhile, compared to PKD1 inhibition by Gȍ6976 treatment alone which is known to alleviate Ras OIS(Fu and Rubin, 2011), the ectopic expression of CBX8-S2E mutant significantly reversed the suppressive effect of Gȍ6976 on Ras OIS (Fig. S6C-J). These results indicate that PKD1 regulates cell senescence at least partly through PKD1-mediated CBX8 S256/311 phosphorylation.

### Phosphorylation of CBX8 at S256/311 by PKD1 impairs PRC1 complex integrity and mitigates H2AK119 mono-ubiquitination

As CBX8 is known to prevent cell senescence via repressing INK4a/ARF locus, and we found that CBX8 S256/311 phosphorylation by PKD1 promotes cell senescence and p16 expression, we then dissected the underlying mechanisms how CBX8 S256/311 phosphorylation affects its function. We first assessed whether PKD1 had impact on CBX8 protein expression in hepatocellular cancer cell line HepG2. HepG2 cells expressed PKD1 and CBX8 (Fig. S7A). Our results showed that either ectopic expression of PKD1 or knockdown of PKD1 by siRNA in a dose-dependent manner did not visibly alter CBX8 protein expressions (Fig. S7B and C). In addition, the kinase activity of PKD1 had no impact on CBX8 expression since either PKD1-CA or PKD1-KD did not alter CBX8 protein expressions (Fig. S7D). These results suggest that PKD1 has no impact on CBX8 protein expression.

Given that CBX8 S256/311 residues are close to the Pc box, which physically interacts with other critical PRC1 components including RING1A/B and BMI1(Chittock et al., 2017; Connelly and Dykhuizen, 2017), we then investigated whether PKD1-mediated CBX8 S256/311 phosphorylation might affect CBX8 binding to other PRC1 components such as its direct binding partner RING1B. Intriguingly, compared to WT-CBX8, we found that the association between phosphomimic mutant CBX8-S2E and RING1B was significantly reduced, whereas the interaction between unphosphorylatable mutant CBX8-S2A and RING1B was significantly increased in HEK293T cells (Fig. 5A). Additionally, the binding between CBX8-S2A and BMI1 was also increased. Moreover, introducing an active PKD1-CA attenuated the association between WT-CBX8 and RING1B as well as BMI1 in cells, while the increased interaction between the unphosphorylatable mutant CBX8-S2A with RING1B and BMI1 remained unchanged either in the absence or presence of PKD1-CA (Fig. 5B), which suggests that PKD1-mediated CBX8 S256/311 phosphorylation is a determining factor for CBX8 interaction with RING1B. Similarly, we also observed that PKD1-CA-mediated CBX8 phosphorylation notably mitigated PRC1 components association in both IMR90 and 2BS cells (Fig. 5C, and Fig. S8A). Furthermore, in both Ras-induced senescent IMR90 cells and replicative senescent 2BS cells in which PKD1 was activated, CBX8-S2A preserved the interaction with RING1B and BMI1, whereas the much weaker binding between WT-CBX8 with RING1B and BMI1 was detected (Fig. 5D, and Fig. S8B). Collectively, these results support the hypothesis that CBX8 phosphorylation at S256/311 by PKD1 reduces the association between CBX8 with RING1B and BMI1 thereby impairing PRC1 complex integrity.

**Figure 5.**
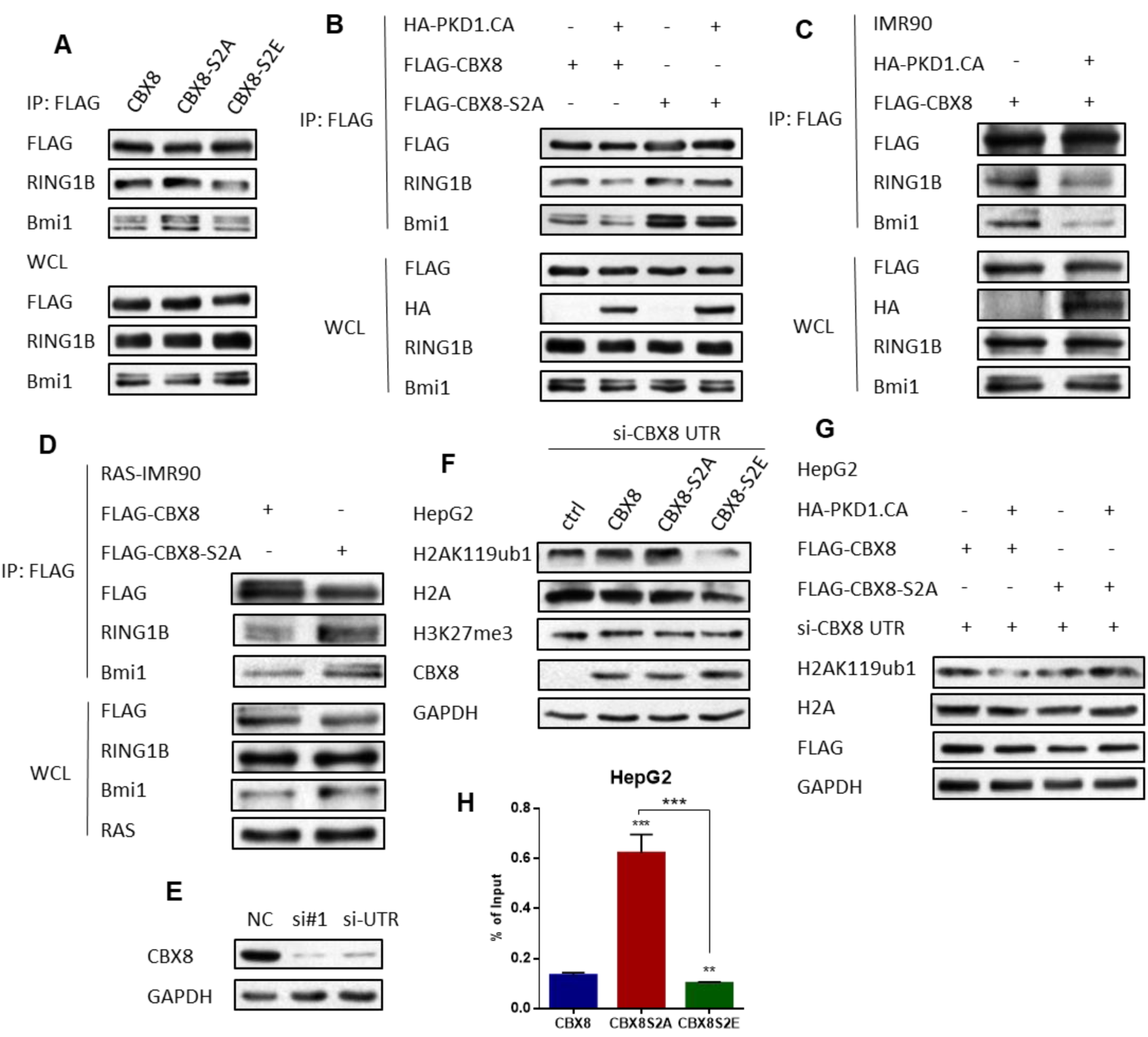
PKD1-mediated CBX8 S256/311 phosphorylation disrupts PRC1 complex integrity and attenuates H2AK119ub1 level. (A) HEK293T cells were transfected with FLAG-WT-CBX8, CBX8-S2A or CBX8-S2E. Then IP assay was carried out using anti-FLAG antibody followed by immunoblotting with the indicated antibodies. (B) HEK293T cells were co-transfected with FLAG-WT-CBX8 or FLAG-CBX8-S2A mutant, with or without HA-PKD1.CA. Then IP assay was carried out using anti-FLAG antibody followed by immunoblotting with the indicated antibodies. (C) IMR90 cells co-transfected FLAG-CBX8 with or without HA-PKD1.CA, then IP assay was carried out using anti-FLAG antibody followed by immunoblotting with the indicated antibodies. (D) RAS-IMR90 cells were transfected with FLAG-WT-CBX8 or FLAG-CBX8-S2A mutant and induced to senescence, then IP assay was carried out using anti-FLAG antibody followed by immunoblotting with the indicated antibodies. (E) HEK 293T cells were transfected with CBX8-siRNA#1 or si-UTR, and CBX8 protein level was detected by western blot. (F) HepG2 cells were transfected with CBX8 UTR siRNA, then infected with FLAG-WT-CBX8, CBX8-S2A mutant or CBX8-S2E mutant. Then cells were subjected to western blot for the indicated proteins. (G) HepG2 cells were transfected with CBX8 UTR siRNA, then infected with FLAG-WT-CBX8 or FLAG-CBX8-S2A mutant, and with or without HA-PKD1.CA. Then cells were subjected to western blot for the indicated proteins. (H) HepG2 cells were transfected with FLAG-WT-CBX8, CBX8-S2A or CBX8-S2E. Then ChIP assay was performed using anti-FLAG antibody to measure the binding activity of WT-CBX8 and its mutants to the INK4A/ARF locus promoter.

Although only RING1A/B are ubiquitin E3 ligases that catalyze the monoubiquitination of H2AK119 (H2AK119ub1) in PRC1 components, however, a stable PRC1 complex is required for its optimal E3 ligase activity(Cao et al., 2005; Elderkin et al., 2007). As PKD1-mediated CBX8 phosphorylation disrupted PRC1 complex integrity, we sought to investigate whether phosphorylation of CBX8 by PKD1 alters H2AK119ub1 level. To this end, we utilized CBX8 3’-UTR siRNA (si-CBX8 UTR) that targeting CBX8 mRNA 3’-UTR to knockdown CBX8 (Fig. 5E), and then re-introduced empty vector, WT-CBX8, CBX8-S2A, or CBX8-S2E plasmids into HepG2 cells, respectively. Compared to the vector control cells, re-expression of WT-CBX8 increased H2AK119ub1 level. Importantly, re-expression of CBX8-S2A mutant further enhanced H2AK119ub1 level relative to WT-CBX8, while re-introduction of CBX8-S2E mutant led to the dramatic loss of H2AK119ub1 level (Fig. 5F). In addition, we noticed that CBX8 S256/311 phosphorylation status had no effect on H3K27me3 level (Fig. 5F). Consistently, in CBX8 knockdown cells, co-expression of WT-CBX8 and PKD1-CA attenuated H2AK119ub1 level compared to CBX8 expression alone. Conversely, co-expression of CBX8-S2A and PKD1-CA did not alter H2AK119ub1 level when compared with CBX8-S2A expression alone (Fig. 5G, and Fig. S8C). Altogether, these result suggest that CBX8 phosphorylation at S256/311 by PKD1 compromises the E3 ligase activity of PRC1 complex and decreases H2AK119ub1 level.

As CBX8 is known to epigenetically repress INK4a/ARF locus transcription, and CBX8 phosphorylation by PKD1 affects p16 expression, we further investigated whether CBX8 S256/311 phosphorylation affects CBX8 binding ability to INK4a/ARF locus promoter in HepG2 cells. Using chromatin immunoprecipitation (ChIP) assay we found that, compared to WT-CBX8, CBX8-S2A mutant largely increased the binding activity to the INK4a/ARF locus, whereas CBX8-S2E mutant reduced that binding (Fig. 5H). Hence, PKD1-mediated CBX8 phosphorylation at S256/311 may lead to CBX8 dissociation from chromatin and thus de-repression of INK4a/ARF locus transcription.

### CBX8 S256/311 phosphorylation attenuates CBX8 oncogenic function in promoting hepatocellular cancer cell proliferation and migration

CBX8 is frequently overexpressed in a wide spectrum of human malignancies including liver, breast, and lymphoma cancers, etc.(Beguelin et al., 2016; Chung et al., 2016; Zhang et al., 2018). Hyper-activation of PRC1/CBX8 leads to epigenetic silencing of various tumor suppressor genes including p16 to facilitate tumor proliferation and metastasis(Bracken et al., 2007; Dietrich et al., 2007). As CBX8 S256/311 phosphorylation compromises PRC1 function and de-represses p16 expression, we further investigated the biological importance of CBX8 S256/311 phosphorylation in tumorigenesis. To this end, we chose two different malignant grade hepatocellular cancer cell lines (HCC) HepG2 and SMMC-7721 to study the functional output of CBX8 S256/311 phosphorylation. Using p-PKD1-S744/S748, p-CBX8-S256, and p-CBX8-S311 antibodies, we found that CBX8 S256/311 phosphorylation level and PKD1 activity were low in more aggressive HepG2 cells, while CBX8 phosphorylation level and PKD1 activity increased in less aggressive SMMC-7721 cells (Fig. 6A). As expected, stable expression of WT-CBX8 in both HepG2 and SMMC-7721 cells suppressed p16 mRNA and protein expression levels relative to empty vector control cells (Fig. 6B, C, and Fig. S9A, B). Notably, compared to WT-CBX8, stable expression of CBX8-S2A was more potent in repressing p16 expression, whereas CBX8-S2E substantially lost the ability to repress p16 expression. The overexpression of WT-CBX8 and CBX8 mutants was confirmed by western blot analysis (Fig. 6B, C, and Fig. S9A, B). As a consequence, compared to WT-CBX8, CBX8-S2A was more potent in promoting HepG2 cell proliferation as evidenced by CCK-8 assay and colony formation assay, whereas CBX8-S2E barely enhanced HepG2 cell proliferation (Fig. 6D and E). Moreover, compared to WT-CBX8, CBX8-S2A was more potent in promoting HCC cells motility and migration as measured by wound healing assay and transwell migration assay, respectively, in both HepG2 and SMMC-7721 cells (Fig. 6F-I, and Fig. S9C-F). On the other hand, CBX8-S2E was unable to augment cancer cell motility and migration (Fig. 6F-I, and Fig. S9C-F). Taken together, these results suggest that CBX8 S256/311 phosphorylation compromises the oncogenic role of CBX8 in hepatocellular cancer cells.

**Figure 6.**
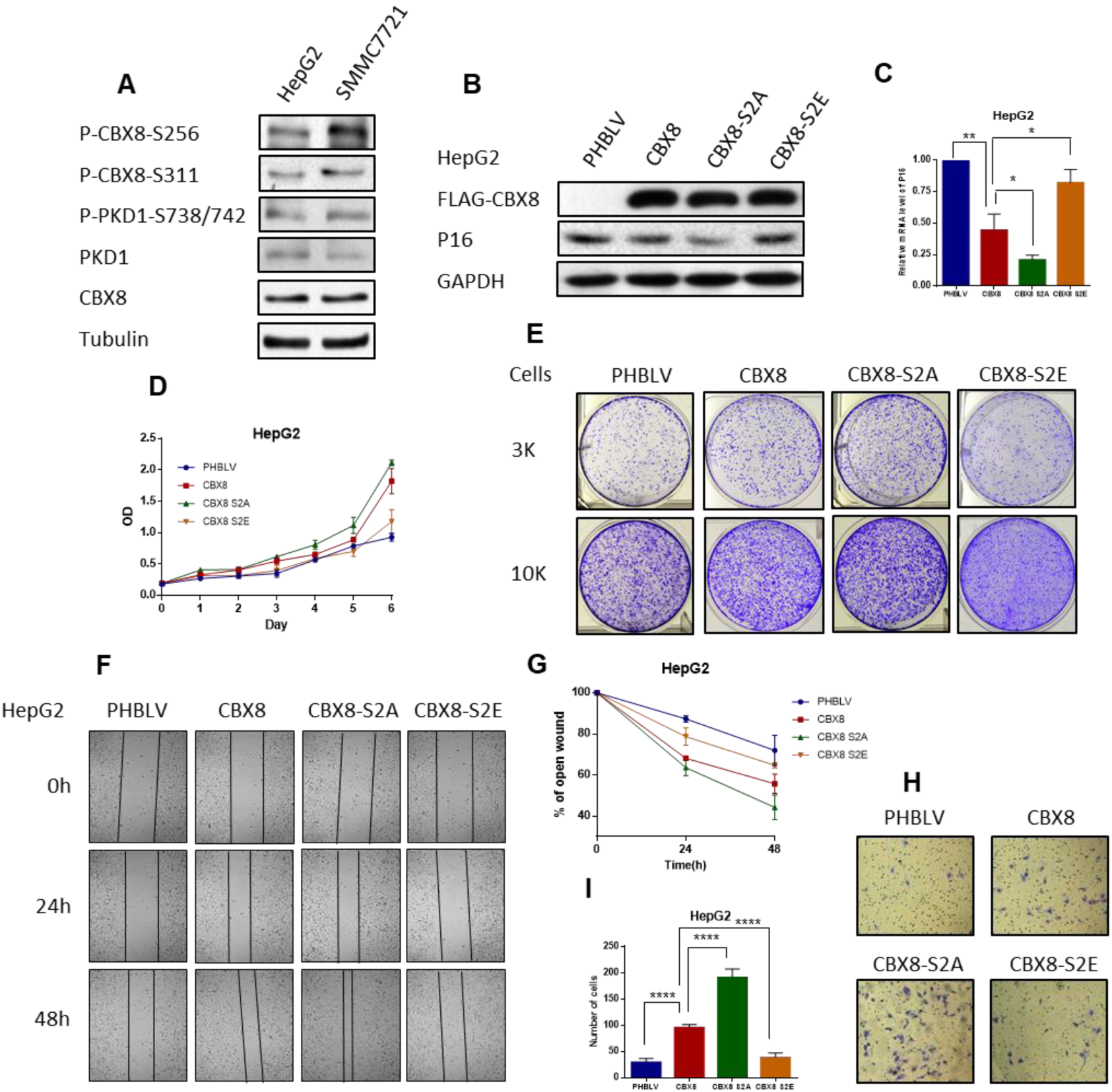
CBX8 S256/311 phosphorylation reduces hepatocellular cancer cells proliferation and migration. (A) Total proteins were extracted from normal HepG2 or SMMC7721 cells, and then subjected to western blot for the indicated proteins. (B) FLAG-WT-CBX8 or FLAG-CBX8-S2A/S2E lentiviral vectors were infected in HepG2 cells, and then cell lysates were subjected to western blot for the indicated proteins. (C) Total RNAs were extracted from (B) cells, then p16 mRNA level was measured by qPCR. (D) Growth curves of HepG2 cells infected with WT-CBX8 or CBX8-S2A/S2E mutants were determined by CCK-8 assay. Data are presented as mean ± s.d. from three independent experiments, each performed in triplicate. (E) HepG2 cells infected with the indicated vectors were cultured for 10 days, then colony formation assay was performed by crystal violet staining cells. (F) HepG2 cells were infected with the indicated vectors. Then artificial wounds were made to cells when cells reach confluence. Images were taken at 0, 24, and 48h after wound, and the wound widths were measured. (G) Data from (F) were quantified and graphed. Error bars represent means ± SD (n = 3). (H) HepG2 cells stably expressing the indicated plasmids were subjected to transwell migration assay, then images were taken and cells were counted. (I) Data from (H) were quantified and graphed. Error bars represent means ± SD (n = 3). ****P < 0.001.

## Discussion

PcG proteins are subjected to the post-translational modifications, particularly phosphorylation, in response to the environment or cell signaling cues (Chen et al., 2010; Kaneko et al., 2010; Kawaguchi et al., 2017; Liu et al., 2012; Voncken et al., 2005; Wan et al., 2018; Wei et al., 2011; Wu et al., 2013; Wu and Zhang, 2011). CBX8 phosphorylation sites have been identified by global phosphoproteomic analyses, yet have never been validated in vivo and the relevant kinases and functions remain totally unknown(Zhou et al., 2013). PIM1 is once reported to phosphorylate CBX8 to promote its degradation thereby up-regulating p16 to enhance cell senescence, however, the phosphorylation site has not been identified in that study. In this study, we have found that PKD1 directly interacts with CBX8 and phosphorylates CBX8 at Ser256 and Ser311. Notably, the motif containing Ser256 and Ser311 are evolutionarily conserved, suggesting that this regulatory mechanism could be functional in multiple organisms (Fig. 1 and 2). Using two antibodies we generated that specifically recognizes CBX8 phosphorylation at Ser256 or Ser311, we further revealed that PKD1 activation triggered either by serum stimulation, or Nocodazole treatment, or Ras OIS, all promotes CBX8 S256/311 phosphorylation (Fig. 3 and Fig. 4A). These results suggest that PKD1-mediated CBX8 S256/311 phosphorylation could be functional in multiple biological processes. To our knowledge, this is the first time to unveil the specific phosphorylation sites of CBX8 in vitro and in vivo and identify its upstream kinase. We further identify that the chromodomain (CD) of CBX8 is critical for CBX8 association with PKD1. Other than CBX8, we also found that PKD1 directly interacts with PRC1 complex component PHC2b, however, PKD1 did not phosphorylates PHC2b and other PRC1 complex components.

CBX8 is well known to suppress cell senescence by epigenetically silencing of INK4A/ARF locus, which encodes p16(Dietrich et al., 2007). CBX8 is down-regulated and dissociates from INK4A/ARF locus in senescent cells, leading to the upregulation of p16(Bracken et al., 2007). We observed that CBX8 is increasingly phosphorylated at S256/311 by PKD1 during the establishment of Ras OIS. We further demonstrate that, compared to WT-CBX8, the unphosphorylatable mutant CBX8-S2A is more potent in repressing cell senescence and p16 expression, whereas the phospho-mimic mutant CBX8-S2E partially compromises the ability to suppress cellular senescence and p16 expression (Fig. 4). These results indicate that PKD1-mediated CBX8 S256/311 phosphorylation facilitates cellular senescence by derepressing p16 expression. Mechanistically, we found that PKD1-mediated CBX8 S256/311 phosphorylation impairs the binding between CBX8 with BMI1 and RING1B thus disrupting the PRC1 complex integrity, which results in the decrease of both H2AK119ub1 level and the binding of CBX8 to p16 promoter (Fig. 5). Both the intact PRC1 and PRC2 complexes are required for efficient epigenetic silencing of target genes(Aranda et al., 2015). Therefore, it is reasonable that the impairment of PRC1 complex integrity caused by PKD1-mediated CBX8 S256/311 phosphorylation ultimately leads to the de-repression of p16 and cellular senescence. These results suggest that PKD1-mediated CBX8 S256/311 phosphorylation is the novel mechanism that PKD1 regulates cell senescence. We previously report that PKD1 activates both NF-κB and classical protein secretory pathway to modulate SASP factors expression and secretion thereby promoting cellular senescence (Su et al., 2018; Wang et al., 2014). Therefore, all of these findings suggest that PKD1 promotes cellular senescence via multiple mechanisms.

Mounting evidences demonstrate that activated PKD1 directly phosphorylates Class-IIa histone deacetylases (HDACs) such as HDAC5 and HDAC7 in the nucleus, leading to their dissociation from the transcription factors and nuclear export, thereby altering histone and chromatin acetylation level and de-repressing the target genes transcription at epigenetic level to regulate diverse biological process (Fielitz et al., 2008; Jensen et al., 2009; Vega et al., 2004; Wang et al., 2008; Xu et al., 2007). However, whether PKD1 can modulate other histone and chromatin modifications such as ubiquitination and methylation remains totally unknown. Here we found that PKD1 directly phosphorylates PRC1 complex component CBX8 at S256/311 thereby impairing PRC1 complex integrity, which leads to the decrease of both H2AK119ub1 level and the binding of CBX8 to p16 promoter, ultimately results in the de-repression of p16 at epigenetic level and cell senescence (Fig. 5). To our knowledge, this is the first time to uncover that PKD1 can directly phosphorylates PcG protein to modulate histone ubiquitination level thereby regulate biological processes at epigenetic level.

CBX8 is aberrantly overexpressed in multiple human cancers including liver, breast, and lymphoma cancers, and CBX8 is known to enhance cancer cells proliferation, migration and invasion(Beguelin et al., 2016; Chung et al., 2016; Zhang et al., 2018). In two different malignant grade HCC cells SMMC-7721 and HepG2, we found that CBX8 S256/311 phosphorylation level and PKD1 activity are low in more aggressive HepG2 cells, while CBX8 phosphorylation level and PKD1 activity increase in less aggressive SMMC-7721 cells. Compared to WT-CBX8, the gain-of-function mutant CBX8-S2A is more potent to promote HCC cells proliferation and migration, whereas the loss-of-function mutant CBX8-S2E mitigates HCC cells proliferation and migration (Fig. 6). PKD1 is reported to promote pancreatic and breast cancer development(Karam et al., 2012; Liou et al., 2015), however, some studies suggest that PKD1 might have anti-cancer function in certain types of cancers including gastric and prostate cancers(Youssef and Ricort, 2019). It is worth to exploit whether PKD1-mediated CBX8 phosphorylation at S256/311 is an important mechanism that PKD1 exerts its anti-cancer function in the future.

In summary, we demonstrate that PKD1 directly interacts with CBX8 and phosphorylates CBX8 at Ser256/311 in several biological processes. PKD1-mediated CBX8 S256/311 phosphorylation impairs PRC1 complex integrity, which leads to the decrease of H2AK119ub1 level and CBX8 dissociation from p16 promoter and subsequent de-repression of p16 and cell senescence. CBX8 S256/311 phosphorylation also attenuates liver cancer cells proliferation and migration.

## Materials and methods

### Cell Culture and Senescence Induction

HEK293T cells, HeLa cells, HepG2 cells, SMMC-7721 cells were cultured in DMEM medium supplemented with 10% FBS. H1299 cells were cultured in RPMI 1640 medium containing 10% FBS. IMR90 human diploid fibroblasts (HDFs) were purchased from the American Type Culture Collection (ATCC) and cultured in DMEM with 10% FBS. ER:Ras-IMR90 cells were generously provided by Masashi Narita, Cancer Research U.K., Cambridge Research Institute, and were given 100nM 4-hydroxytamoxifen (4-OHT) to induce ER-Ras fusion protein expression and maintained in 4-OHT–containing DMEM (w/o phenol red) with 10% FBS until harvesting. Human diploid 2BS fibroblasts (National Institute of Biological Products, Beijing, China) were cultured in RPMI 1640 supplemented with 10% FBS.

### Antibodies and Reagents

The following primary antibodies were used for immunoblotting analysis: anti-PKD1 (90039), CBX8 (14696), HA (3724), p-MOTIF PKD1 substrate (4381), anti-phosphothreonine (9381), p-PKD1 (Ser744/748) (2054), H3K27me3 (9733), H2AK119ub (8240), p-H3 (9849), cyclin B1 (12231) were purchased from Cell Signaling Technology (CST); anti-phosphoserine (ab9332), anti-H2A (ab177308), H3 (ab1791), Bmi1 (ab126783), RING1B (ab181140) were purchased from Abcam; anti-GST(sc-965), anti-RAS (sc-32), anti-IgG (sc-2025) were from Santa Cruz; anti-p16 (10883-1-AP), anti-p21 (60214-1-Ig) were from Proteintech; anti-tubulin (BS1482M, BioWorld); anti-FLAG (F3165, Sigma); anti-p-CBX8 (Ser256) generated by Biodragon; anti-p-CBX8 (Ser311) generated by Genscript.

DMSO (Sigma) was used as a solvent to dissolve PMA (Sigma) and Gö6976 (Sigma). Nocodazole (Selleck) and 4-OHT (Sigma) were dissolved in methanol.

### Immunoprecipitation, silver staining and mass spectrometry

Cells were collected and lysed in IP lysis buffer (25mM Tris-HCl (pH 7.4), 150 mM NaCl, 1% NP-40, 1mM EDTA, and 5% glycerol) mixing with protease inhibitor cocktail (Sigma) at 4°C for 30 min. The lysates were incubated with protein A/G Agarose beads for 2 h at 4 °C to pre-clean. The extracts were incubated with FLAG antibody or mouse IgG overnight at 4 °C, then incubated with the protein A/G Agarose beads for 2 h at 4 °C. The beads were washed five times with lysis buffer. The immunoprecipitates were dissolved in 2×SDS loading buffer and resolved on NuPAGE 4–12% Bis-Tris gel (Invitrogen), and then silver stained using Pierce silver stain kit (Thermo). The protein bands were retrieved and subjected to LC–MS/MS analysis.

### GST pull-down assay and in vitro kinase assay

GST and GST-tagged protein were expressed in BL21 (DE3) cells and affinity-purified with glutathione Sepharose 4B affinity chromatography according to the manufacture instructions. FLAG-PKD1 and FLAG-PKD1-CA protein were expressed in HEK293T cells and purified with anti-FLAG affinity Beads (SMART) in accordance with the manufacture instructions. The purified FLAG-PKD1 (500 ng) and GST or GST tagged protein (500 ng/each) were incubated together in 500 μl BC100 buffer at 4 °C overnight. Glutathione-sepharose beads (GE Healthcare) were added and incubated for 2-4 h at 4°C. The beads were washed five times with BC100 buffer. The reaction mixture was boiled in Laemmli buffer. Western blotting was performed using antibody against FLAG and GST.

For in vitro kinase assay, 2 μg of fusion proteins and 8 μg of FLAG-PKD1-CA were incubated in kinase buffer (Cell Signaling Technology) for 30 min at 30°C in the presence of 200 μM ATP. Then SDS loading buffer was added to stop the reaction. The phosphorylated proteins were analyzed by Western blotting with anti-phosphoserine, anti-phosphothreonine, or anti-pMOTIF PKD1 substrate antibodies.

### Immunoblotting

Cell pellets were lysed in RIPA buffer (Applygen Technologies) containing phosphatase inhibitor (Roche Diagnostics) and protease inhibitor (Fermentas). Cell lysates were then centrifuged for 15 min at 15,000 g at 4°C, and the insoluble debris was discarded. Protein concentration was measured using the BCA Protein Assay Kit (Pierce). Cell lysates (20-40 μg) were subjected to 8-15% SDS-PAGE and transferred to nitrocellulose membranes (Millipore). The membrane was blocked using 5% milk in TBST buffer at room temperature for 1 h. Primary antibodies were blotted using 5% milk or BSA in TBST, and incubated at 4 °C overnight. The HRP-conjugated anti-mouse or anti-rabbit secondary antibodies were incubated for 1 h at room temperature in 5% milk/TBST. Then the signals were detected by enhanced chemiluminescence ECL (Pierce, Thermo Scientific), and imaged by films.

### Real-time PCR

Total RNA was extracted using the RNeasy Mini kit (Qiagen) following the manufacturer’s protocol and then subjected to reverse transcription using the StarScript first strand cDNA synthesis kit (Transgen Biotech, Beijing, China). Real-time PCR was performed using SYBR Select Master Mix (Applied Biosystems) on an ABI PRISM 7500 Sequence Detector (Applied Biosystems). GAPDH served as an internal control for normalization.

The primers for RT-qPCR are listed as below: p16 forward: 5′-CGGTCGGAGGCCGATCCAG-3′ and reverse: 5′-GCGCCGTGGAGCAGCAGCAGCT-3′ GAPDH forward: 5′-CGACCACTTTGTCAAGCTCA -3′ and reverse: 5′-AGGGGTCTACATGGCAACTG -3′

### Plasmids and siRNAs Transfection

Full-length CBX8 and CBX8-S2A or S2E mutant plasmids were transiently transfected into cells with PEI Reagent following the manufacturer’s instruction. Two independent siRNA sequences against CBX8 were: siRNA#1: 5′-CUCGCUUGCUCGCAGCCUU -3′, and siRNA UTR: 5′-GCGUGAGCUUGGCAUAGUG -3′. Two independent siRNA sequences against PKD1 were: siRNA#1: 5′-GUCGAGAGAAGAGGUCAAAUU -3′, and siRNA UTR: 5′-CAAUCCUCAUUGUUUCGAAAU -3′. siRNAs were transfected using Lipofectamine RNAiMAX Reagent following the manufacturer’s protocol. After 72 h transfection, cells were harvested and lysed to evaluate the transfection efficiency.

### Lentiviral Vectors and Viral Infection

pHBLV-CBX8 and pHBLV-CBX8-S2A or S2E mutant vectors were constructed by cloning the full-length CBX8 and CBX8-S2A or S2E mutant fragments into the EcoRI/XbalI sites of pHBLV-puro vector. Lentiviral constructs were transfected with packaging plasmids to HEK293T cells. The supernatant containing lentiviral particles was harvested at 48 h and filtered through a 0.45 μm filter, and then directly added to the culture medium. The infected cells were then selected with puromycin.

### SA-β-Gal activity assay

SA-β-gal staining was performed according to the manufacturer’s protocol (Beyotime, Shanghai, China). Cells were washed twice with ice-cold 1×PBS, and fixed in 3% formaldehyde for 10 min at room temperature. After washing twice with 1×PBS, cells were stained at 37°C overnight in SA-β-gal staining solution without CO2, then images were acquired by photomicrography. 500 cells were counted to calculate the percentages of cells positive for SA-β-gal staining.

### EdU incorporation assay

To perform EdU incorporation assay, 1 × 10^4^ cells were cultured in 96-well plate and performed assay with Cell-Light EdU Apollo567 In Vitro Kit (C10310-1, Ribobio) according to the manufacturer’s instructions. All cells were examined using fluorescence microscopy (Leica) with the appropriate filters. At least 500 cells were counted in randomly chosen fields from each culture well for statistical analysis.

### Colony formation assay

To perform colony formation, 3 × 10^3^, and 1 × 10^4^ cells were cultured in six-well plate. After 10-14 days, cells were fixed in 4% (wt/vol) formaldehyde at 37 °C for 30 min and washed twice with 1× PBS, then stained with crystal violet for 1h and washed twice with 1× PBS followed by photography.

### Cell Growth Curves

Cell proliferation was detected using 2-(2-Methoxy-4-nitrophenyl)-3-(4-nitrophenyl)-5-(2,4-disulfophenyl)-2H-tetrazoliumsodiumsalt (CCK-8/WST-8) method. 2 × 103 cells per well were seeded into 96-well plate and cultured for periods ranging from 1 to 6 day. The medium was changed every 24 h. At the indicated times, an aliquot of cells was stained with 10 μL of CCK-8 solution (Dojindo) for 2 h, and then the optical density at 450 nm was determined.

### Wound-healing assay

To perform wound-healing assay, cells were seeded in confluent monolayers in six-well plates after transfection. A linear gap was generated by scratching the cell layer at the bottoms of the wells using a sterile 200 uL pipette tip. Phase contrast images were acquired at an identical location at 0, 24 and 48 h after scratching, and the width (W) of the scratch wound was measured. The rate of closure of the open wounds was calculated. All measurements were performed in triplicate at least three times.

### Transwell migration assay

Double-chamber migration assay was performed using transwell chambers (24-well plate, 8um pores; BD Biosciences). In brief, the lower chambers were filled with 750uL DMEM containing 10% FBS. HCC cells with different treatments were suspended in serum-free medium, seeded in the upper chambers and incubated at 37°C for 24 h. Then, the cells on the upper surface of the filters were removed using cotton wool swabs. The migrated cells on the lower side of the membrane were fixed in 95% methanol and stained with 0.1% crystal violet dye, and the number of cells migrating to the lower surface was counted in three randomly selected high-magnification fields (100×) for each sample.

### Serum Starvation and Serum Induction

HEK 293T cells or H1299 cells were subjected to serum starvation (standard medium + 0% FBS) for 24 h, then serum was added (standard medium + 10% FBS), and cells were harvested at various time points.

### Statistical Analysis

Two-tailed unpaired Student’s t-test was used to determine the significance of differences between samples indicated in figures. Results are depicted as mean values ± standard deviation (SD, n=3). P < 0.05 (*), P < 0.01 (**), P < 0.005 (***), or P < 0.001 (****) were considered significant.

## Acknowledgements

This work was supported by grants from Ministry of Science and Technology of the People’s Republic of China (2018YFC2000102), and from National Natural Science Foundation of China (81571369 and 31871382).

## Author Contributions

Y.Y.S. and J.C. designed research; Y.Y.S., L.Y., C.Z.X., N.Z, D.D.L and G.D.L. performed the experiments and analyzed the data; T.J.T supervised the research; Y.Y.S., and J.C. wrote the paper.

