## supplemental information for "Protein kinase D1 phosphorylates CBX8 to facilitate the disassociation of PRC1 complex from p16 promoter and promotes cell senescence"

Jun Chen

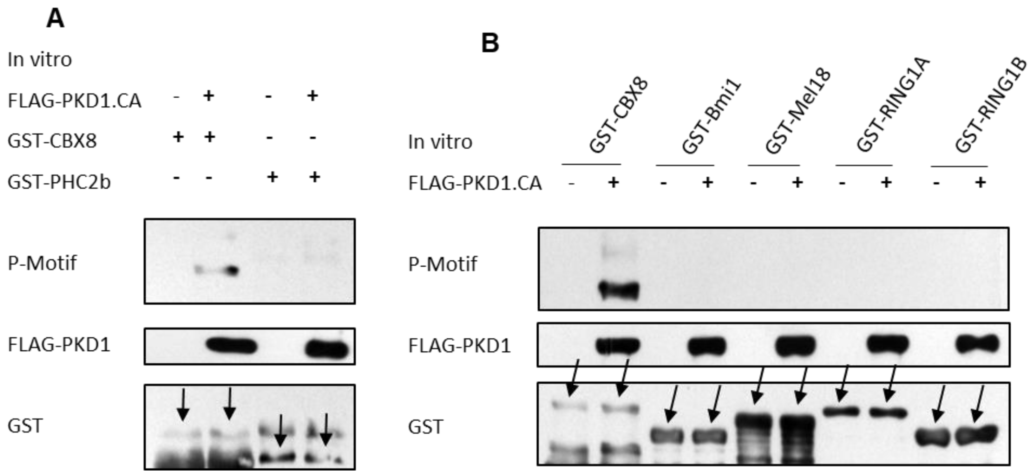


**Figure S1. PKD1 only phosphorylates CBX8, but not other PRC1 complex components (related to Figure 1)**

(A) Purified GST-tagged CBX8 or PHC2b proteins were incubated with or without FLAG-PKD1.CA in the presence of ATP. The reaction products were separated by SDS-PAGE and immunoblotted with the indicated antibodies.

(B) Purified GST-tagged CBX8, Bmi1, Mel18, RING1A or RING1B proteins were incubated with or without FLAG-PKD1.CA in the presence of ATP. The reaction products were separated by SDS-PAGE and immunoblotted with the indicated antibodies.

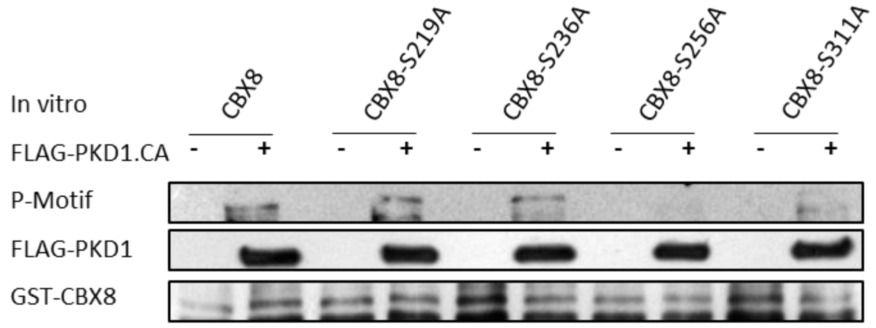


**Figure S2. PKD1 phosphorylates CBX8 at Ser256 and Ser311 (related to Figure 2)**

(A) Purified GST-tagged WT-CBX8 or its various mutant proteins were incubated with or without FLAG-PKD1.CA in the presence of ATP. The reaction products were separated by SDS-PAGE and immunoblotted with the indicated antibodies.

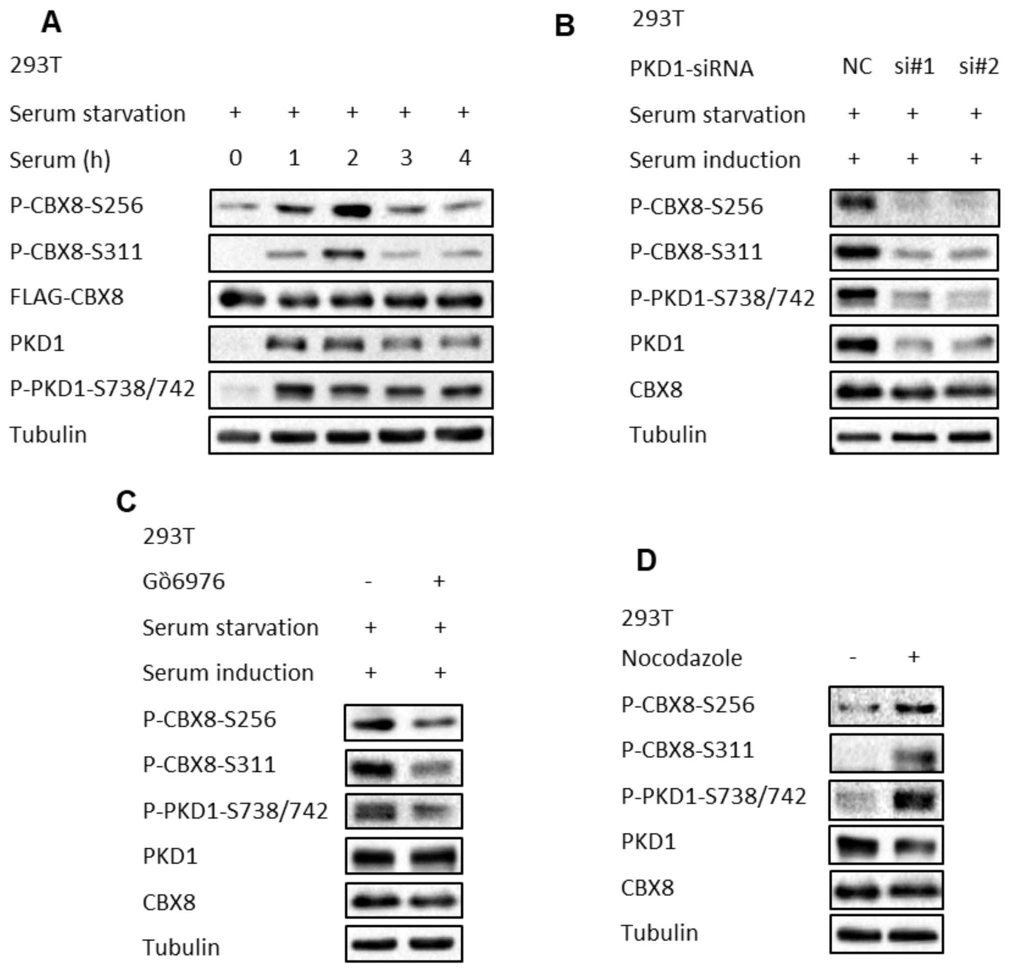


**Figure S3. Activation of PKD1 by serum stimulation or Nocodazole treatment promotes CBX8 S256/311 phosphorylation (related to Figure 3)**

(A) 293T cells were deprived of serum for 24h, then serum was reintroduced in the culture for the indicated times. Then cells were collected and subjected to western blot for the indicated proteins.

(B) 293T cells were transfected with PKD1-siRNAs for 48 h, then serum was deprived for 24h. Cells were then added serum for 1h and harvested for immunoblotting with the indicated antibodies.

(C) 293T cells were deprived of serum for 24h with or without Gò6976 treatment, then serum was reintroduced for 1h. Then cells were collected and subjected to western blot for the indicated proteins.

(D) 293T cells were treated with or without Nocodazole (100 ng/mL) for 16h, then cells were subjected to immunoblotting with the indicated antibodies.

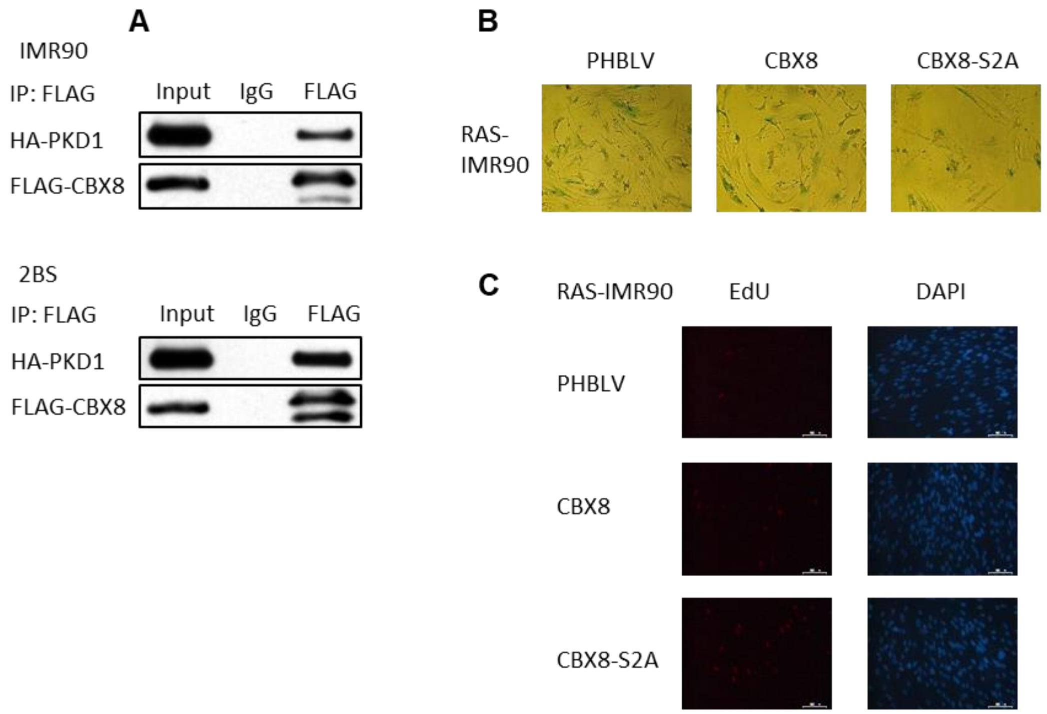


**Figure S4. Unphosphorylatable mutant CBX-S2A is more potent than WT-CBX8 to suppress Ras OIS (related to Figure 4)**

(A) IMR90 or 2BS cells were transfected with FLAG-CBX8 and HA-PKD1 plasmids. IP assay and subsequent western blot analysis were performed by using FLAG and HA antibodies.

(B) RAS-IMR90 cells infected with the indicated vectors were subjected to SA-β-gal activity staining. The representative images were shown.

(C) RAS-IMR90 cells infected with the indicated vectors were subjected to EdU incorporation assay. The representative images of EdU incorporation (red) and DAPI staining (blue) were shown.

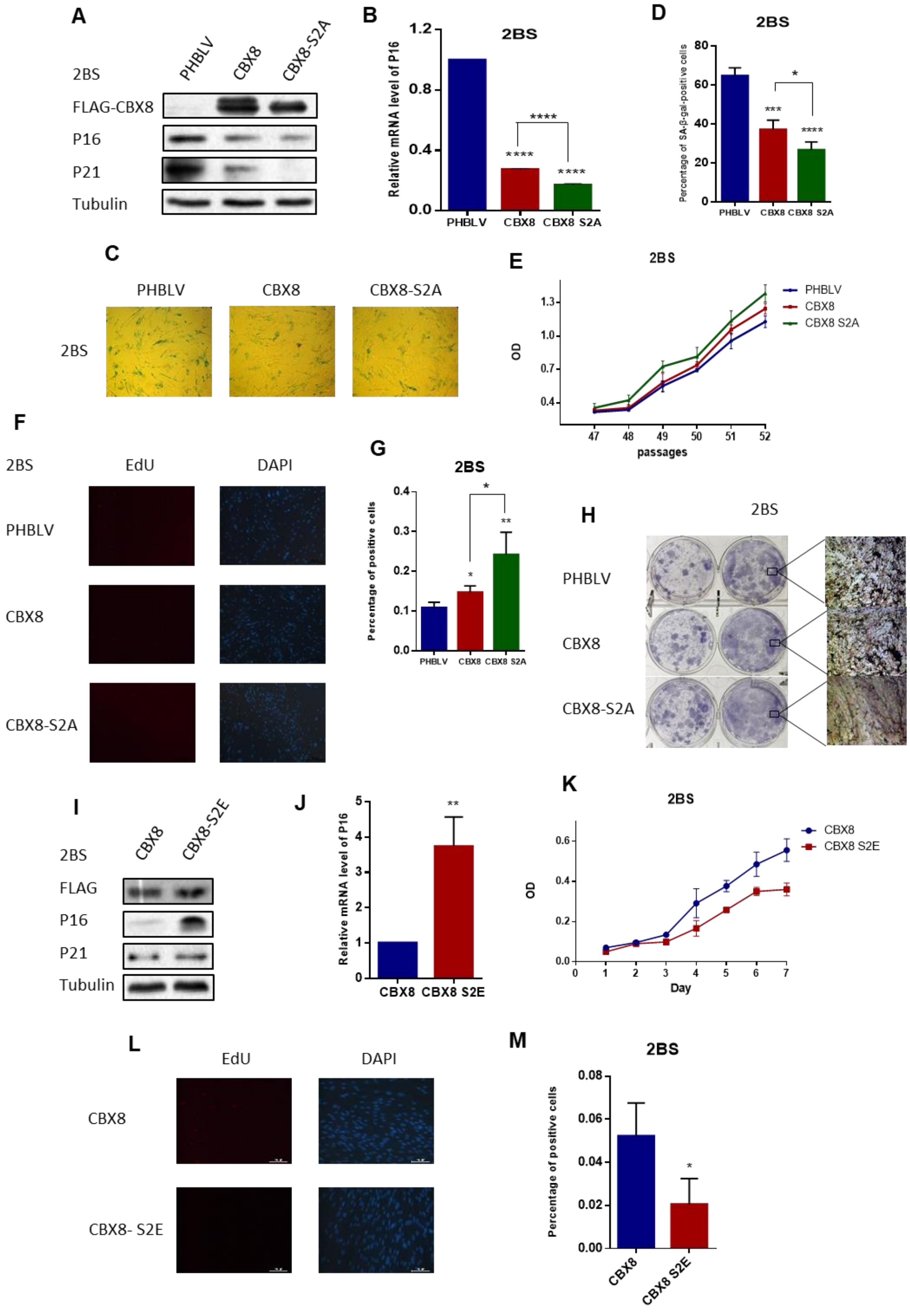


**Figure S5. CBX8 S256/311 phosphorylation status affects 2BS cells replicative senescence (related to Figure 4)**

(A) 2BS cells were infected with the indicated lentiviral vectors. Then cells were subjected to western blot for the indicated proteins.

(B) Total RNAs were extracted from (A) cells, then p16 mRNA level was measured by qPCR.

(C) 2BS cells infected with the indicated vectors were subjected to SA-β-gal activity staining. The representative images were shown.

(D) The percentages of SA-β-gal staining positive cells were statistically analyzed and graphed. Data are presented as mean ± s.d. from three independent experiments with 300 cells per experiment.

(E) Growth curves of 2BS cells infected with the indicated vectors were determined by CCK-8 assay. Data are presented as mean ± s.d. from three independent experiments, each performed in triplicate.

(F) 2BS cells infected with the indicated vectors were subjected to EdU incorporation assay. The representative images of EdU incorporation (red) and DAPI staining (blue) were shown.

(G) The percentages of EdU incorporating positive cells were statistically analyzed and graphed. Data are presented as mean ± s.d. from three independent experiments with 300 cells per experiment.

(H) 2BS cells infected with the indicated vectors were cultured for 14 days, then colony formation assay was performed by crystal violet staining cells. The representative magnified images (x40) were shown.

(I-M) 2BS cells were infected with FLAG-WT-CBX8 or FLAG-CBX8-S2E mutant lentiviral vectors. Then cells were subjected to western blot (I), realtime qPCR (J), cell growth assay (K), EdU incorporation assay (L and M), to evaluate cell senescent status.

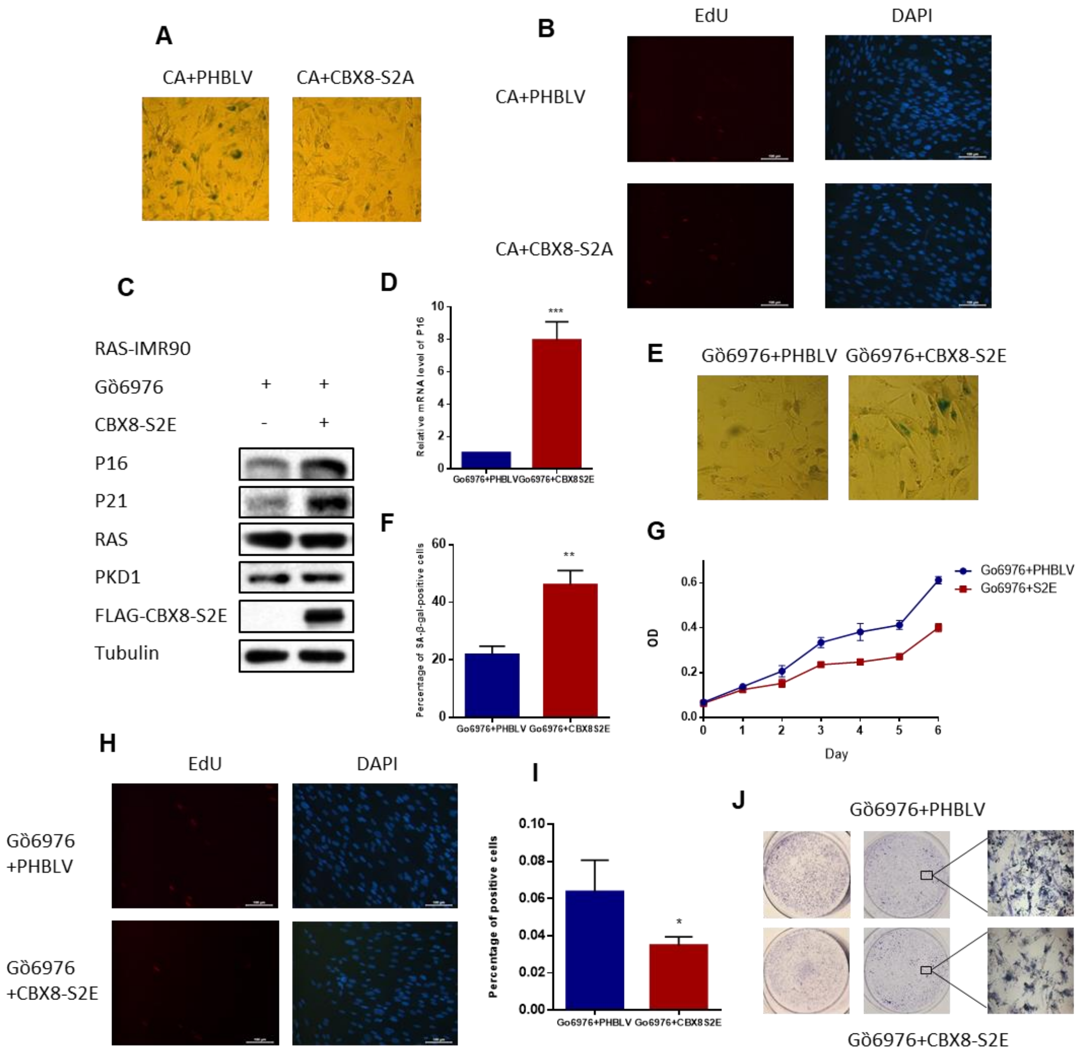


**Figure S6. PKD1 regulates Ras OIS partly through PKD1-mediated CBX8S256/311 phosphorylation (related to Figure 4)**

(A and B) RAS-IMR90 cells were infected with HA-PKD1.CA lentiviral vector, then transfected with or without CBX8-S2A. Cell were subjected to SA-β-gal activity staining (A) or EdU incorporating assay (B). The representative images were shown.

(C-J) RAS-IMR90 cells were infected with or without CBX8-S2E, then treated with Gö6976. Cell were subjected to western blot (C), realtime qPCR (D), SA-β-gal activity staining (E and F), cell growth assay (G), EdU incorporating assay (H and I), colony formation assay (J), to evaluate cell senescent status.

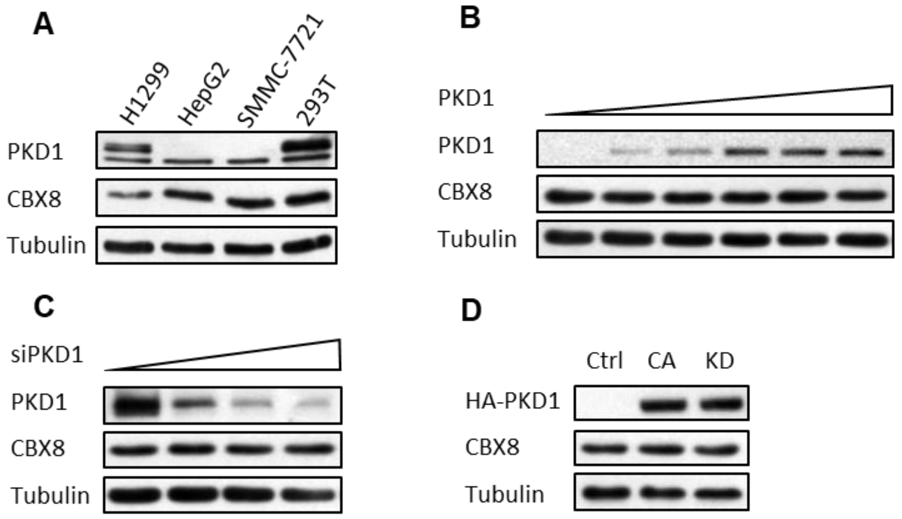


**Figure S7. PKD1 does not alter CBX8 protein expression**

(A) H1299, HepG2, SMMC-7721, and HEK293T cells were subjected to western blot analysis for the indicated proteins.

(B) HepG2 cells were transfected with increasing amount of HA-PKD1 plasmid, and then subjected to western blot analysis for the indicated proteins.

(C) H1299 cells were transfected with increasing amount of PKD1-siRNA, and then subjected to western blot analysis for the indicated proteins.

(D) HEK 293T cells were transfected with HA-PKD1.CA or KD, and then immunoblotted with the indicated antibodies.

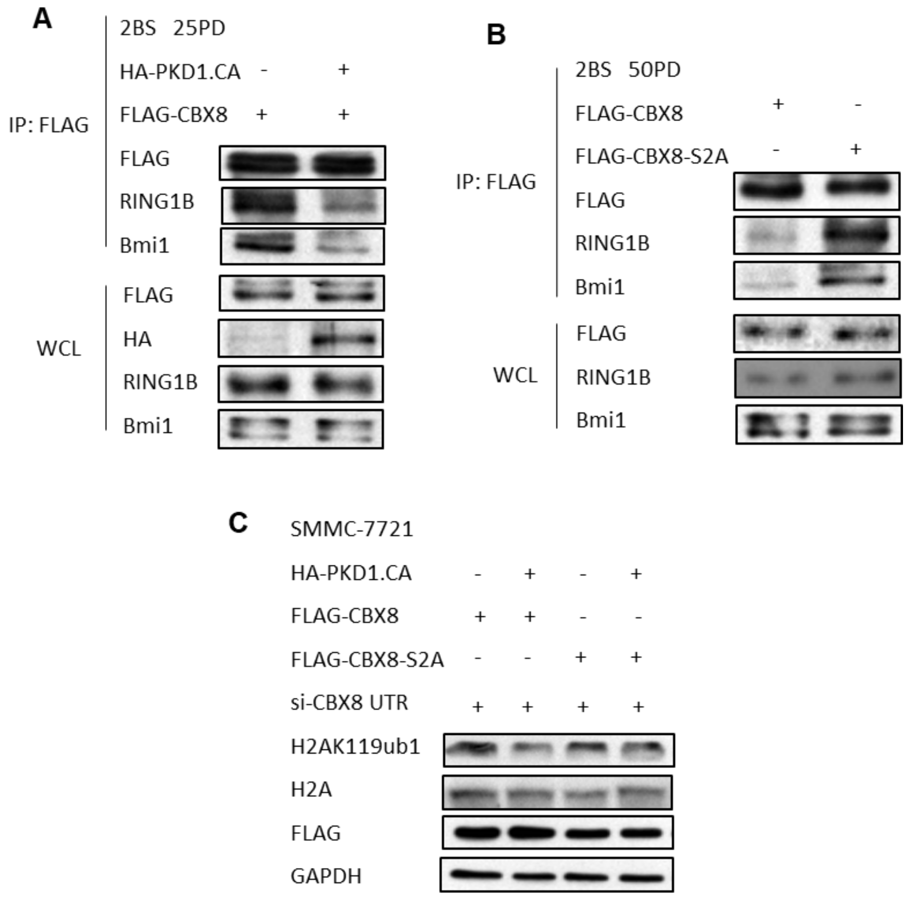


**Figure S8. PKD1-mediated CBX8 S256/311 phosphorylation impairs PRC1 complex integrity and attenuates H2AK119ub1 level (related to Figure 5)**

(A) 25PD 2BS cells co-transfected FLAG-CBX8 with or without HA-PKD1.CA, then IP assay was carried out using anti-FLAG antibody followed by immunoblotting with the indicated antibodies.

(B) 50PD 2BS cells transfected with FLAG-WT-CBX8 or FLAG-CBX8-S2A mutant, then IP assay was carried out using anti-FLAG antibody followed by immunoblotting with the indicated antibodies.

(C) SMMC-7721 cells were transfected with CBX8 UTR siRNA, then infected with FLAG-WT-CBX8 or FLAG-CBX8-S2A mutant, and with or without HA-PKD1.CA. Then cells were subjected to western blot for the indicated proteins.

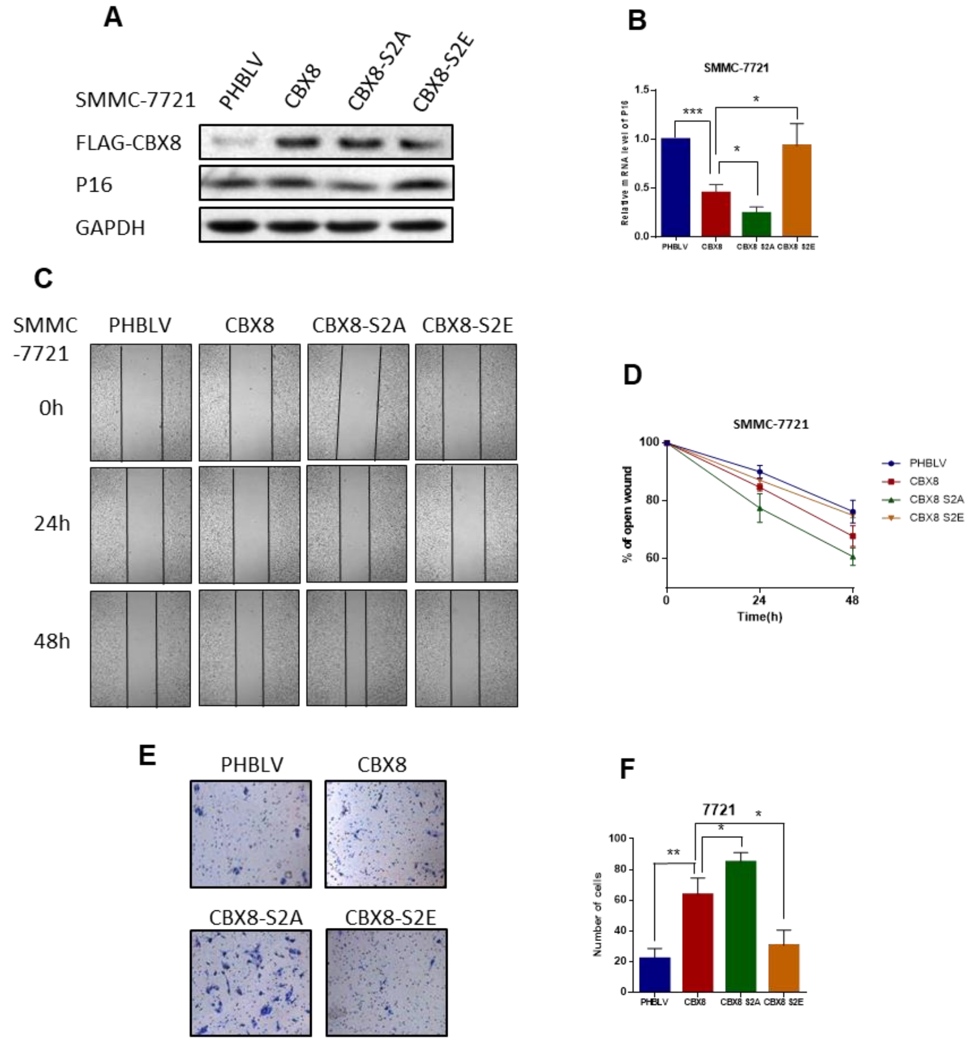


**Figure S9. CBX8 S256/311 phosphorylation decreases HCC cells proliferation and migration (related to Figure 6)**

(A) FLAG-WT-CBX8 or FLAG-CBX8-S2A/S2E lentiviral vectors were infected in SMMC-7721 cells, and then cell lysates were subjected to western blot for the indicated proteins.

(B) Total RNAs were extracted from (A) cells, then p16 mRNA level was measured by qPCR.

(C) SMMC-7721 cells were infected with the indicated vectors. Then artificial wounds were made to cells when cells reach confluence. Images were taken at 0, 24, and 48h after wound, and the wound widths were measured.

(D) Data from (C) were quantified and graphed. Error bars represent means ± SD (n = 3).

(E) SMMC-7721 cells stably expressing the indicated plasmids were subjected to transwell migration assay, then images were taken and cells were counted.

(F) Data from (E) were quantified and graphed. Error bars represent means ± SD (n = 3). *P < 0.05, **P < 0.01.
